# Systems genetics identifies ETS1 as a stress-dependent regulator of adipocyte insulin action and heme–iron homeostasis

**DOI:** 10.64898/2026.09.01.748508

**Authors:** Yi Lin Jiang, Kristen C. Cooke, Harry B. Cutler, Meg Potter, Alexis Diaz-Vegas, Stewart W. C. Masson, Marin E. Nelson, Jonathan Scavuzzo, Aaron Lambert, James G. Burchfield, Gregory J. Cooney, Grant Morahan, Jacqueline Stöckli, David E. James, Søren Madsen

**Author notes:** These authors contributed equally (Yi Lin Jiang, Kristen C. Cooke).

## Abstract

White adipose tissue plays a central role in systemic energy homeostasis by buffering nutrient excess through insulin-stimulated glucose uptake and triglyceride storage. Despite its importance, the genetic and molecular mechanisms governing adipose tissue insulin action remain poorly defined because tissue-specific insulin responsiveness has been difficult to quantify at the scale required for genetic discovery. Here, we developed the first scalable platform for high-throughput genetic mapping of tissue-specific insulin action in adipose tissue, enabling systems genetic analysis across 559 genetically diverse Diversity Outbred Australia (DOz) mice. Genetic analysis accounting for adiposity identified 39 loci associated with adipose tissue insulin action, demonstrating that adipose insulin responsiveness is a genetically encoded trait that captures a dimension of metabolic health beyond adiposity. Among these, a strong diet-dependent locus on chromosome 9 encompassed the transcription factor *Ets1*. Functional studies demonstrated that *Ets1* silencing selectively restored insulin-stimulated glucose uptake in insulin-resistant adipocytes. Proteomic profiling revealed that ETS1 orchestrates a stress-responsive program involving heme metabolism, iron handling and redox homeostasis. Consistent with this, ETS1 knockdown reduced cellular heme and labile iron levels and attenuated oxidative stress under insulin-resistant conditions. Collectively, these findings demonstrate the power of systems genetics to identify previously unrecognised regulators of adipose insulin action and establish the heme–iron axis as a critical determinant of adipocyte insulin responsiveness.

## Introduction

White adipose tissue (WAT) is now recognised as a central regulator of systemic metabolic homeostasis^1^. Traditionally viewed as a passive reservoir for excess energy, WAT protects other organs from nutrient overload by storing excess calories as triglycerides, thereby limiting ectopic lipid accumulation and lipotoxicity^1,2^. More recently, however, adipose tissue has emerged as a dynamic endocrine organ that communicates extensively with other tissues through the secretion of adipokines^1,3^. These include leptin, which regulates appetite and energy expenditure via the central nervous system, and adiponectin, which promotes insulin sensitivity and lipid homeostasis in peripheral tissues^3,4^. These discoveries have fundamentally changed our understanding of adipose tissue from a simple energy store to a major regulator of whole-body metabolism.

A third, and equally important, function of adipose tissue is to act as an insulin-responsive nutrient sensor, coupling glucose flux into adipocytes to whole-body metabolic homeostasis^5^. Indeed, adipocytes and skeletal muscle have served as the archetypal model systems for understanding insulin-regulated glucose transport, culminating in the discovery of the insulin-responsive glucose transporter GLUT4^6^. Insulin-stimulated glucose uptake fulfils several essential functions within adipocytes. It supplies glycerol-3-phosphate required for triglyceride synthesis, provides carbon for de novo lipogenesis, and fuels a range of metabolic pathways that contribute to adipocyte function^5^. Although adipose tissue accounts for a smaller proportion of acute whole-body glucose disposal than skeletal muscle, the substantial mass of adipose tissue, particularly in obesity, suggests that it may represent an important long-term sink for circulating glucose^7^. Furthermore, a significant proportion of glucose taken up by adipocytes is converted to lactate and released into the circulation^5,8^, where it may contribute to both hepatic gluconeogenesis and inter-organ metabolic signalling. Together, these observations position adipocyte glucose uptake as a fundamental metabolic process extending well beyond lipid storage.

Multiple lines of evidence indicate that adipocyte insulin action plays an important role in maintaining systemic metabolic health. Adipose-specific deletion of GLUT4 causes whole-body insulin resistance and glucose intolerance despite preserved GLUT4 expression in skeletal muscle^9^, demonstrating that impaired glucose uptake in adipocytes alone is sufficient to disrupt systemic glucose homeostasis. Likewise, human genetic studies have identified variants associated with whole-body insulin resistance that implicate adipose tissue gene regulation^10,11^, including variants influencing GLUT4 expression^12^. Collectively, these findings indicate that adipocyte insulin sensitivity is not merely a consequence of systemic metabolic dysfunction but may itself represent an important determinant of metabolic health.

Despite its physiological importance, little is known about why adipocyte insulin sensitivity varies so substantially between individuals. This gap is particularly relevant to obesity, where individuals with similar degrees of adiposity can exhibit markedly different metabolic phenotypes, a distinction commonly framed as metabolically healthy versus metabolically unhealthy obesity^13,14^. While adipose tissue expandability, inflammation and fibrosis have all been implicated in this heterogeneity^15,16^, adipocyte insulin sensitivity is likely to represent a fundamental component of adipose tissue function not captured by adiposity alone^17,18^. Our previous studies using a diverse panel of inbred mouse strains demonstrated marked gene-by-environment interactions in adipose tissue insulin sensitivity^19^, suggesting that naturally occurring genetic variation is a major determinant of this phenotype. However, the molecular mechanisms responsible remain largely unknown.

Here, we leveraged the extensive natural genetic diversity of the Diversity Outbred Australia (DOz) mouse population^20^ to identify genetic regulators of adipocyte insulin-stimulated glucose uptake. By combining high-resolution genetic analysis with functional studies in adipocytes, we identify the transcription factor ETS1 as a diet-dependent regulator of adipose tissue insulin action. Mechanistically, we demonstrate that ETS1 promotes insulin resistance by disrupting cellular iron homeostasis, increasing oxidative stress, and impairing insulin-stimulated glucose uptake, revealing a previously unrecognised pathway regulating adipocyte insulin sensitivity.

## Results

### Direct phenotyping enables genetic dissection of adipose insulin action

To derive direct measures of adipose insulin action for genetic mapping, we developed a scalable *ex vivo* assay compatible with the genetically heterogeneous DOz mouse resource (Supp. Fig. 1). The assay could directly quantify basal and insulin-stimulated glucose uptake in WAT explants across a range of insulin concentrations, using a 2-deoxy-glucose (2DG) tracer. By measuring insulin responsiveness *ex vivo*, this approach avoids confounding physiological variables including blood flow^21^, circulating hormones^22–24^ and substrate availability^25,26^, while preserving tissue architecture and enabling standardised insulin stimulation across large cohorts.

**Figure 1.**
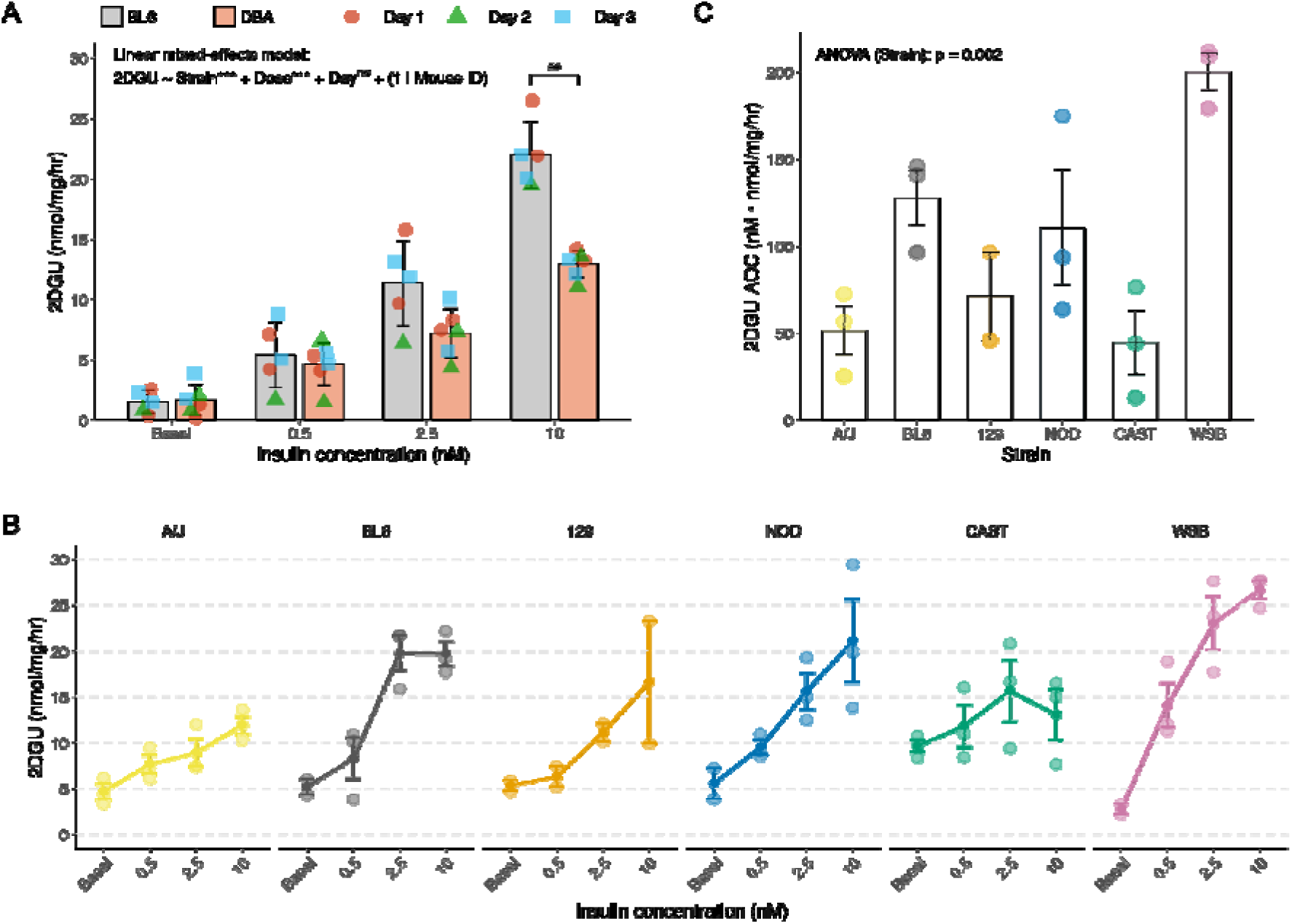
A high-throughput *ex vivo* 2DGU assay reveals strain-specific differences in WAT insulin action. **(A)** Basal and insulin-stimulated 2-deoxyglucose uptake (2DGU) (0.5, 2.5, and 10 nM) in gonadal white adipose tissue (gWAT) from chow-fed C57BL/6J (BL6) and DBA/2J (DBA) mice measured across three consecutive days. A linear mixed-effects model was fitted with strain, insulin dose, and experimental day as fixed effects, and mouse ID as a random intercept. Asterisks denote the statistical significance of each variable (***p ≤ 0.001; ns, not significant). Pairwise comparisons between strains at each dose were assessed by t-test with Benjamini–Hochberg correction (^##^p.adj ≤ 0.01). **(B)** 2DGU measures in gWAT across various inbred mouse strains: A/J, BL6, 129X1/SvJ (129), NOD/ShiLtJ (NOD), CAST/EiJ (CAST), and WSB/EiJ (WSB). **(C)** Area of the curve (AOC) for each mouse strain, calculated from the 2DGU response. Statistical significance of strain effect was assessed by one-way ANOVA.

First, we established that the assay could robustly resolve differences in insulin-stimulated 2DG uptake (2DGU) in gonadal white adipose tissue (gWAT) from mice of different genetic backgrounds, using C57BL/6J (BL6) and DBA/2J (DBA) mice (Fig. 1A). We next extended this analysis to additional inbred strains including founder strains of the DOz population (Fig. 1B). Basal and insulin-stimulated 2DGU varied markedly between strains, with approximately 2-fold differences in basal uptake and substantial differences in the magnitude of the insulin response. WSB/EiJ (WSB) mice exhibited a 9-fold increase in 2DGU following insulin stimulation, whereas CAST/EiJ (CAST) and A/J strains showed more modest 2-fold responses. Notably, basal 2DGU in CAST mice was comparable to the maximal insulin-stimulated response in A/J mice, indicating that both basal glucose uptake and insulin responsiveness are strongly influenced by genetic background. To capture variation across the entire insulin dose-response relationship, we calculated the area of the curve (AOC), integrating both the magnitude and dose-responsive kinetics of the insulin response (Fig. 1C). Together, these data established the first scalable platform for high-throughput genetic mapping of tissue-specific insulin action. The marked strain-dependent variation further suggests that adipose tissue insulin responsiveness is under substantial genetic control.

### Diet and genetics generate extensive variation in adipose tissue insulin action

The DOz population captures extensive natural genetic variation, with each mouse carrying a unique mosaic of founder alleles (Extended Data Fig. 1A-B). Extensive recombination between founder genomes enables high-resolution mapping of complex traits. To identify novel regulators of WAT insulin sensitivity, 559 DOz mice were fed either a chow or WD for 8 or 14 wks prior to metabolic phenotyping and SNP-based genotyping (Fig. 2A). Insulin-stimulated glucose uptake was quantified in gWAT from these mice using our *ex vivo* 2DGU assay.

**Figure 2.**
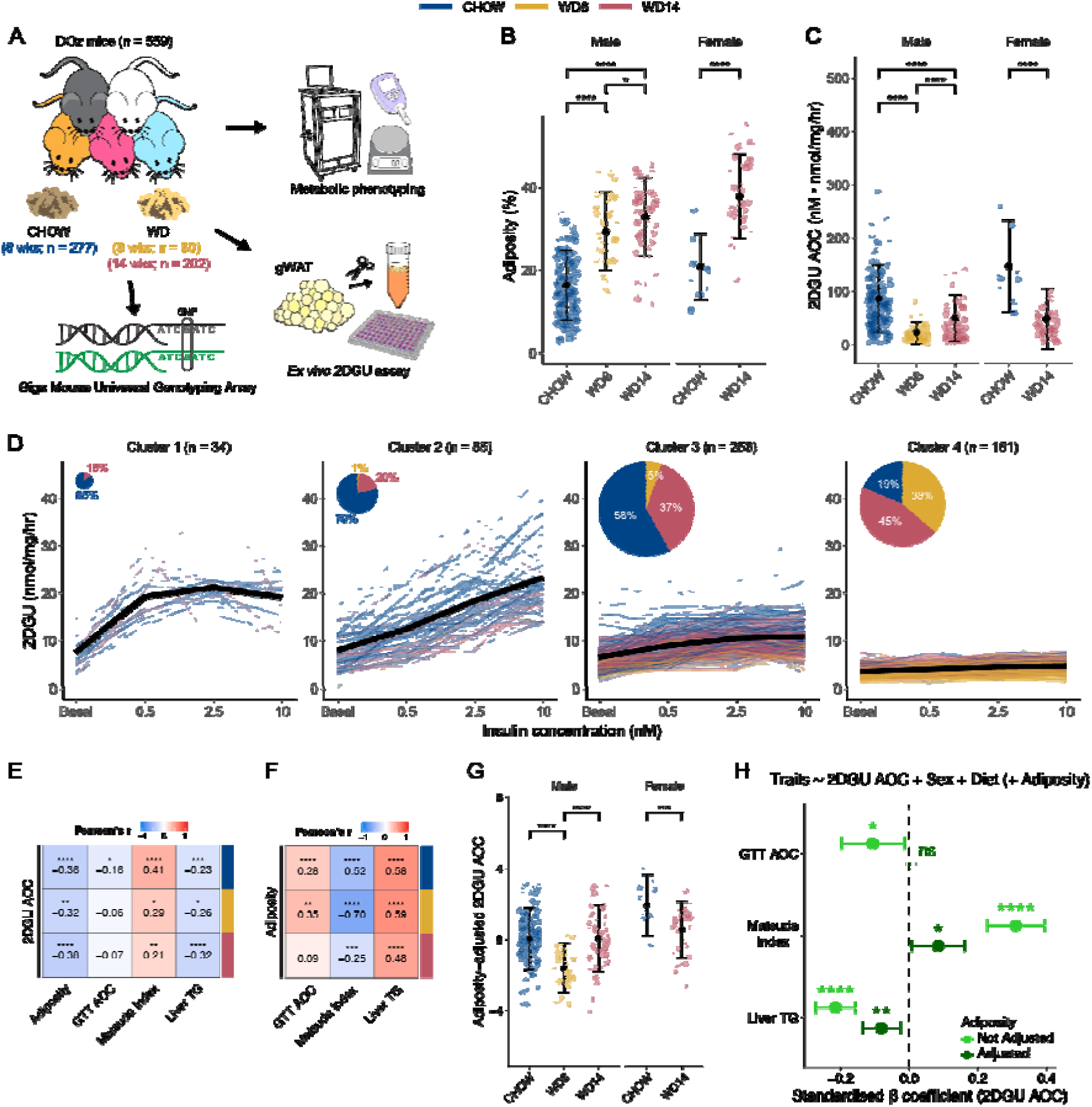
*Ex vivo* 2DGU profiling reveals marked heterogeneity in WAT insulin action in DOz mice. **(A)** Study design: DOz mice were fed either chow or a Western diet (WD) for 8 (WD8) or 14 wks (WD14). SNP, Single Nucleotide Polymorphism. **(B-C)** Adiposity (B) and WAT insulin sensitivity (2DGU area of the curve (AOC)) (C), in DOz mice across sex and diet groups. Data are mean ± SD. Asterisks denote the statistical significance of mean differences between groups: t-test applied for (B), and Wilcoxon test applied for (C) (Benjamini-Hochberg adjustment; *p.adj ≤ 0.05, ****p.adj ≤ 0.0001). **(D)** Hierarchical clustering of individual 2DGU responses identifies four distinct phenotypic groups. Each line represents an individual mouse; colours indicate diet. Pie charts denote cluster sizes and diet distributions. **(E)** Correlations between 2DGU AOC and systemic metabolic traits: adiposity, glucose tolerance test (GTT) AOC, Matsuda Index, and liver triglycerides (TG), stratified by diet. **(F)** Correlations between adiposity and other systemic metabolic traits: GTT AOC, Matsuda Index, and liver TG, stratified by diet. For (E-F), numbers indicate correlation coefficients; asterisks denote statistical significance (Benjamini-Hochberg adjustment; *p.adj ≤ 0.05, **p.adj ≤ 0.01, ***p.adj ≤ 0.001, ****p.adj ≤ 0.0001). **(G)** Adiposity-adjusted 2DGU AOC in DOz mice across sex and diet groups. Adiposity-adjusted 2DGU AOC was calculated as the residuals from a linear regression model with 2DGU AOC as the dependent variable and adiposity as the independent variable. Data are mean ± SD. Asterisks denote the statistical significance of mean differences between groups (t-test; Benjamini-Hochberg adjustment; ***p.adj ≤ 0.001, **** p.adj ≤ 0.0001). **(H)** A linear model was fitted to determine 2DGU AOC associations with GTT AOC, Matsuda Index, and liver TG content, with or without adjusting for adiposity. Points represent standardised regression coefficients (β), and error bars denote 95% confidence intervals. Asterisks denote the statistical significance of each variable (Benjamini-Hochberg adjustment; *p.adj ≤ 0.05, **p.adj ≤ 0.01, ****p.adj ≤ 0.0001; ns, not significant). For (E-H), original adiposity values and Box-Cox-transformed 2DGU AOC, GTT AOC, Matsuda Index, and liver TG values were used.

As expected, WD feeding increased adiposity and induced changes to systemic metabolism (Fig. 2B; Extended Data Fig. 2A-H). However, substantial phenotypic variation remained within dietary groups, with considerable overlap between chow- and WD-fed animals across multiple metabolic traits, highlighting a strong contribution of genetic background to metabolic responses.

Direct measurement of insulin-stimulated 2DGU further revealed pronounced heterogeneity in adipose tissue insulin responsiveness. Although WD feeding reduced WAT insulin responsiveness overall (Fig. 2C), the responses were highly heterogeneous, with many WD-fed animals retaining insulin responsiveness comparable to that of chow-fed mice. Intriguingly, insulin-stimulated 2DGU was on average higher in WD14 than in WD8 (Fig. 2C), likely due to adaptive changes in adipose tissue^27,28^, such as adipocyte hyperplasia, in WD14 but not in WD8 mice. Hierarchical clustering of complete insulin dose-response profiles identified four distinct response states spanning a continuum from highly insulin-responsive (Fig. 2D, left) to functionally unresponsive adipose tissue (Fig. 2D, right; Extended Data Fig. 3A-B). Notably, diet alone did not determine cluster membership, demonstrating substantial inter-individual variation in susceptibility to diet-induced adipose tissue insulin resistance.

We next examined how variation in WAT insulin-stimulated 2DGU related to conventional measures of systemic metabolism. As expected, WAT 2DGU was negatively associated with adiposity and liver triglyceride content, and positively associated with whole-body insulin sensitivity (Matsuda Index), consistent with the *ex vivo* assay capturing physiologically meaningful variation in adipose tissue insulin action (Fig. 2E). However, adiposity was itself significantly associated with multiple systemic metabolic traits (Fig. 2F), raising the possibility that some of the observed variation in WAT 2DGU simply reflected differences in obesity rather than intrinsic differences in adipocyte insulin action.

This distinction was particularly important for the subsequent genetic analyses. Because adiposity is itself a highly heritable trait^29,30^, genetic mapping of WAT 2DGU without accounting for adiposity could identify loci associating with WAT 2DGU through effects on adiposity rather than directly regulating adipocyte insulin action. To focus on genetic determinants of WAT insulin sensitivity independent of obesity, we subsequently adjusted for adiposity when performing genetic mapping on WAT 2DGU traits. Importantly, considerable variation in WAT insulin action remained after this correction (Fig. 2G). Moreover, WAT 2DGU remained significantly associated with liver triglyceride content and the Matsuda Index after adiposity adjustment (Fig. 2H), demonstrating that this residual variation (Fig. 2G) retains physiological relevance. Thus, our strategy was not to remove biologically meaningful variation but rather to separate the genetic regulation of adiposity from that of adipocyte insulin action. After adjusting for adiposity, diet, and sex, measures of WAT insulin action exhibited a narrow-sense heritability (h²) of 0.3–0.4 (Extended Data Fig. 3C), further supporting the use of these covariates in subsequent genetic analysis of adipose tissue insulin action.

### Systems genetics identifies loci associated with adipose tissue insulin action

We performed haplotype- and SNP-based quantitative trait locus (QTL) mapping^31^ and identified 14 haplotype-based QTL and 27 SNP-based QTL associated with the six 2DGU traits (Fig. 3A; Extended Data Table 1). Several loci contained genes previously implicated in glucose metabolism, insulin action and adipocyte biology, such as *Pkn2* and *Lmo4* on chromosome 3^32,33^ and *Slc2a12* on chromosome 10^34,35^ (Fig. 3B, C). These findings support the biological validity of our mapping strategy while identifying numerous previously unrecognised candidate regulators of adipose tissue insulin action.

**Figure 3.**
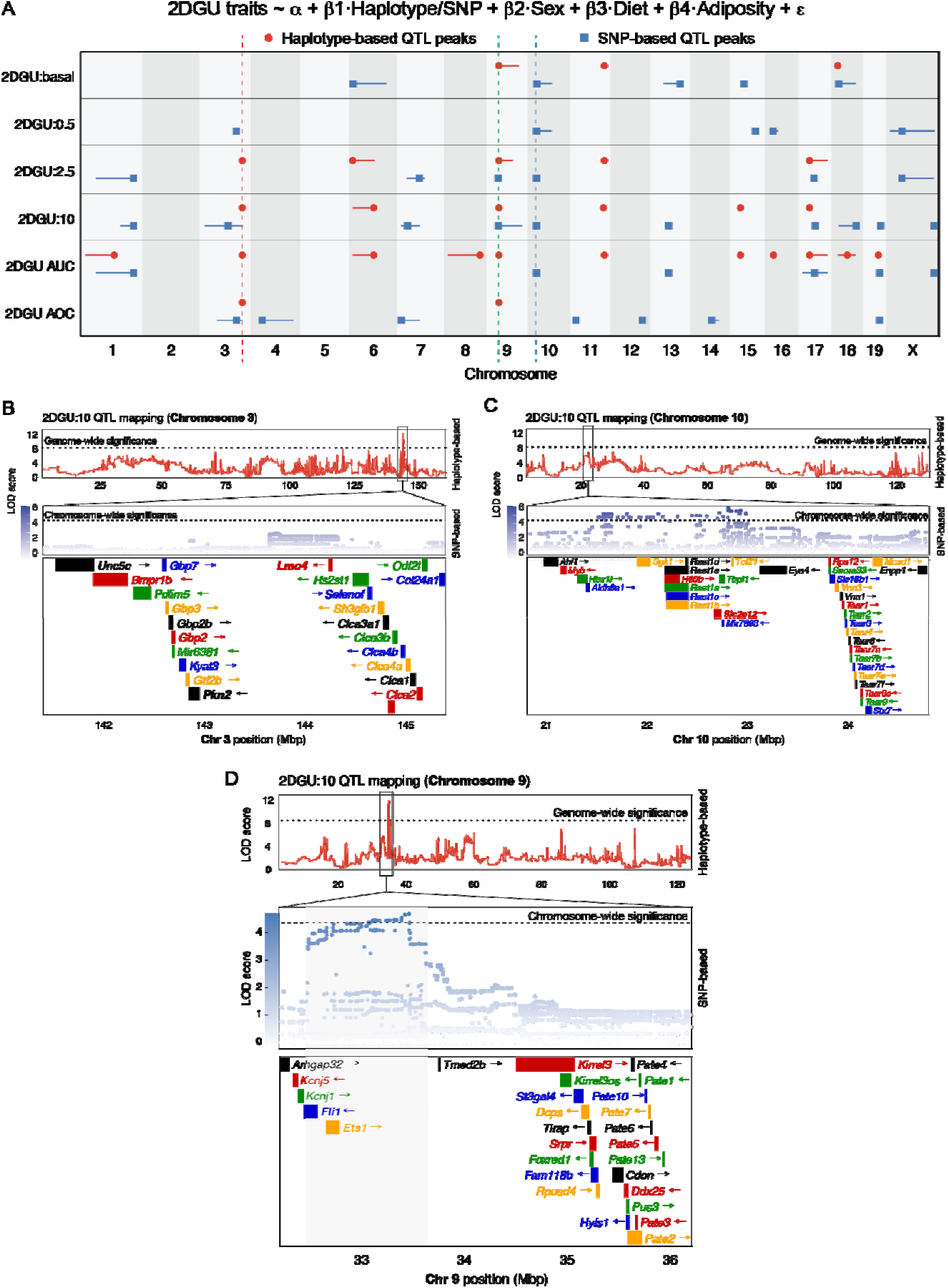
QTL mapping identifies genetic loci associated with variation in WAT insulin action. **(A)** Genome-wide haplotype-(red circles) and SNP-based (blue squares) QTL mapping for basal and insulin-stimulated 2DGU traits (basal or insulin dose indicated), as well as AUC (Area Under the Curve) and AOC calculated from the basal and insulin-stimulated 2DGU measures. Confidence intervals are shown for each QTL peak. Shared haplotype (Chr 3), shared SNP (Chr 10), and the co-localising (Chr 9) loci are indicated by red, blue, and green dashed lines, respectively. The linear mixed model used for mapping is shown schematically. **(B–D)** LOD scores for 2DGU at 10 nM insulin at key loci: shared haplotype peak (Chr 3) (B), shared SNP peak (Chr 10) (C), and the co-localising peak (Chr 9) (D). Haplotype-based results are shown across entire chromosomes with genome-wide significance thresholds. SNP-based associations are shown within ±2 Mb of the peaks, with chromosome-wide significance thresholds. Gene annotations are included.

A particularly compelling signal emerged on chromosome 9 (∼32–36 Mb), where five haplotype-based and two SNP-based QTL converged across multiple 2DGU traits (Fig. 3A, green dashed line; Fig. 3D). Collectively, these analyses identified a resource of 39 loci regulating adipose tissue insulin responsiveness. Among these, the chromosome 9 locus emerged as particularly compelling because of its convergence across multiple mapping strategies, making it the highest-priority locus for mechanistic investigation.

### The genetic variant on chromosome 9 drives enhanced 2DGU

We next focused on the chromosome 9 locus and examined the phenotypic consequences of variation at the lead genotyped SNP by stratifying DOz mice according to their genotype (Fig. 4A). The minor allele (CT or CC genotype) exhibited an allele dosage-dependent effect on insulin-stimulated 2DGU, with animals carrying two copies of the minor allele (CC genotype) showing approximately 20% greater 2DGU at the maximum insulin dose. Strikingly, this effect was observed exclusively in WD-fed mice (Fig. 4A). Moreover, in WD-fed mice, the minor allele was associated with reduced liver triglyceride levels (Fig. 4E), but not with adiposity, glucose tolerance, or whole-body insulin sensitivity (Fig. 4B-D). This pattern validated our adiposity-adjusted mapping strategy as we identified a genetic variant that enhanced insulin-stimulated 2DGU independently of adiposity. Furthermore, these findings support a direct relationship between adipose tissue insulin action and hepatic lipid accumulation^36^.

**Figure 4.**
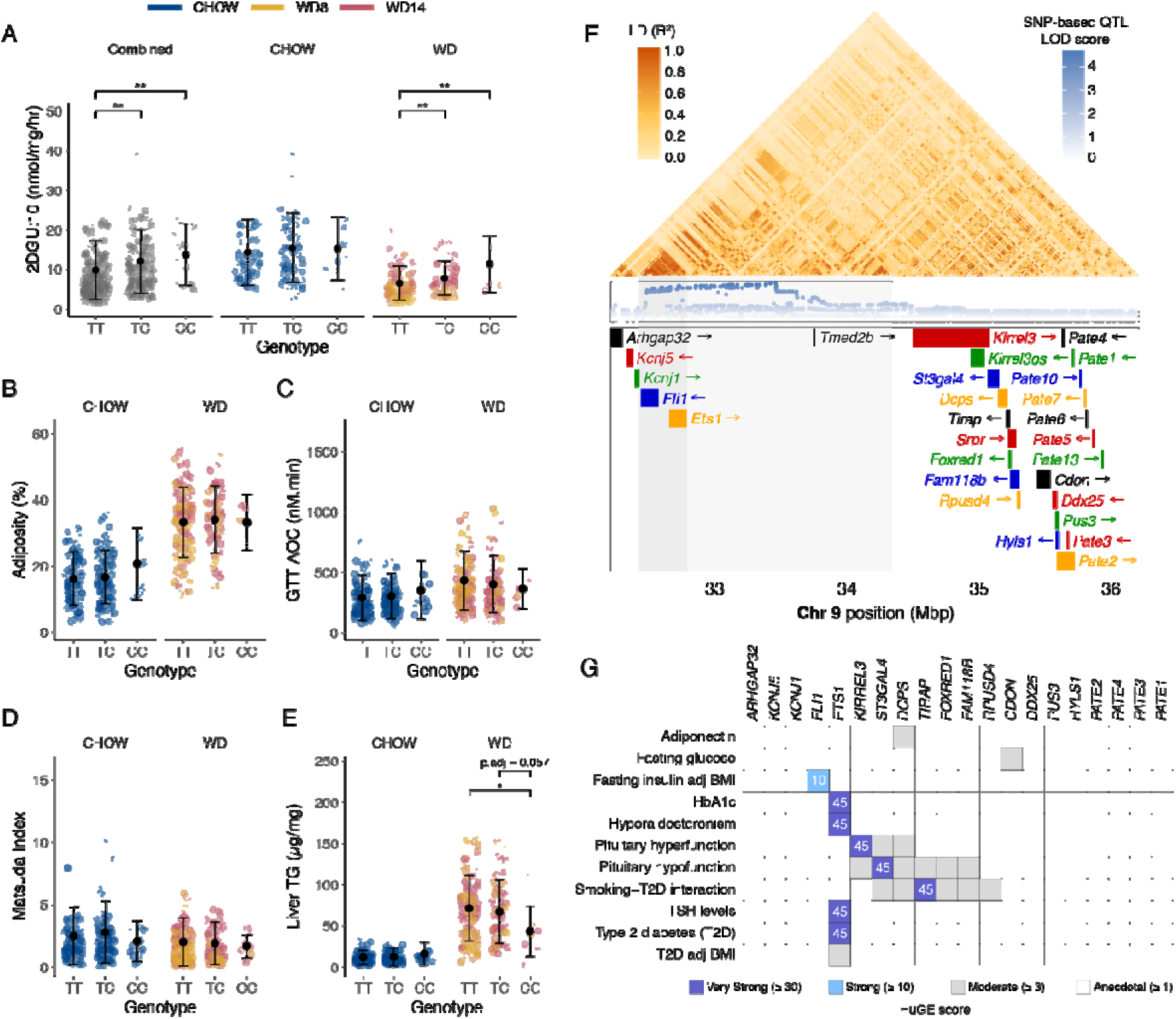
Integration of mouse QTL mapping with human genetics implicates *Ets1* as a diet-dependent regulator of WAT glucose uptake. **(A)** 2DGU at 10 nM insulin across genotypes at the lead SNP in the chromosome 9 locus, shown for all DOz mice and stratified by diet. **(B-E)** Systemic metabolic measures: adiposity (B), glucose tolerance test (GTT) AOC (C), Matsuda Index (D), and liver triglycerides (E), across genotype groups at the lead true SNP within the chromosome 9 locus, stratified by diet. For (A-E), data are mean ± SD; asterisks denote the statistical significance of mean differences between groups (Wilcoxon test; Benjamini–Hochberg correction; *p.adj ≤ 0.05, **p.adj ≤ 0.01). **(F)** Linkage disequilibrium (LD) analysis of SNPs surrounding the co-localising QTL on chromosome 9. The broader LD block containing the high-LOD-score SNPs is highlighted in light grey, and the sub-LD block area containing *Ets1* and *Fli1* is highlighted in dark grey. **(G)** Human Genetics Evidence (HuGE) scores for selected glycaemic traits of human orthologous genes located within the co-localising QTL interval (±2 Mb). HuGE scores ≥ 10 are labelled. TSH, thyroid-stimulating hormone.

### *Ets1* is a candidate regulator of adipose tissue insulin action

To prioritise candidate genes within the chromosome 9 locus, we next integrated genetic, haplotype and human genetic evidence. SNP-based analysis highlighted a smaller region within the QTL with high-LOD-score SNPs (Fig. 3D; highlighted in grey), containing two genes: E26 avian leukemia oncogene 1 (*Ets1*) and Friend leukemia integration 1 (*Fli1*), both of which encode transcription factors. Neither gene has previously been implicated in WAT insulin action.

To refine the underlying genetic interval, we next performed linkage disequilibrium (LD) analysis (Fig. 4F). This identified a 2 Mb region spanning ∼32.5–34.5 Mb around the high-LOD-score SNPs, with a tighter sub-block (R² > ∼0.8) centred on *Ets1* and *Fli1*, indicating that variants within this region likely co-segregate with the lead 2DGU-associated SNPs.

Lastly, we evaluated human genetic support for all genes in the region using Human Genetic Evidence (HuGE) scores via our recently developed web-based tool *Synteny*^37^. Among all genes within the locus, *ETS1* showed the strongest support, with four “Very Strong” associations with human glycaemic traits, including HbA1c and type 2 diabetes (Fig. 4G), whereas *FLI1* showed comparatively little evidence of association.

Together, these complementary analyses prioritised *Ets1* as the strongest candidate causal gene within the chromosome 9 locus, leading us to hypothesise that it functions as a diet-dependent regulator of adipocyte insulin action.

### ETS1 modulates adipocyte insulin action during insulin resistance

To test the above prediction, we silenced *Ets1* in 3T3-L1 adipocytes using retrovirally delivered shRNA (shETS1), with a non-targeting hairpin (shNT) serving as the control. This approach achieved ∼80% knockdown (KD) in ETS1 protein abundance across multiple independent retroviral infections (Fig. 5A).

**Figure 5.**
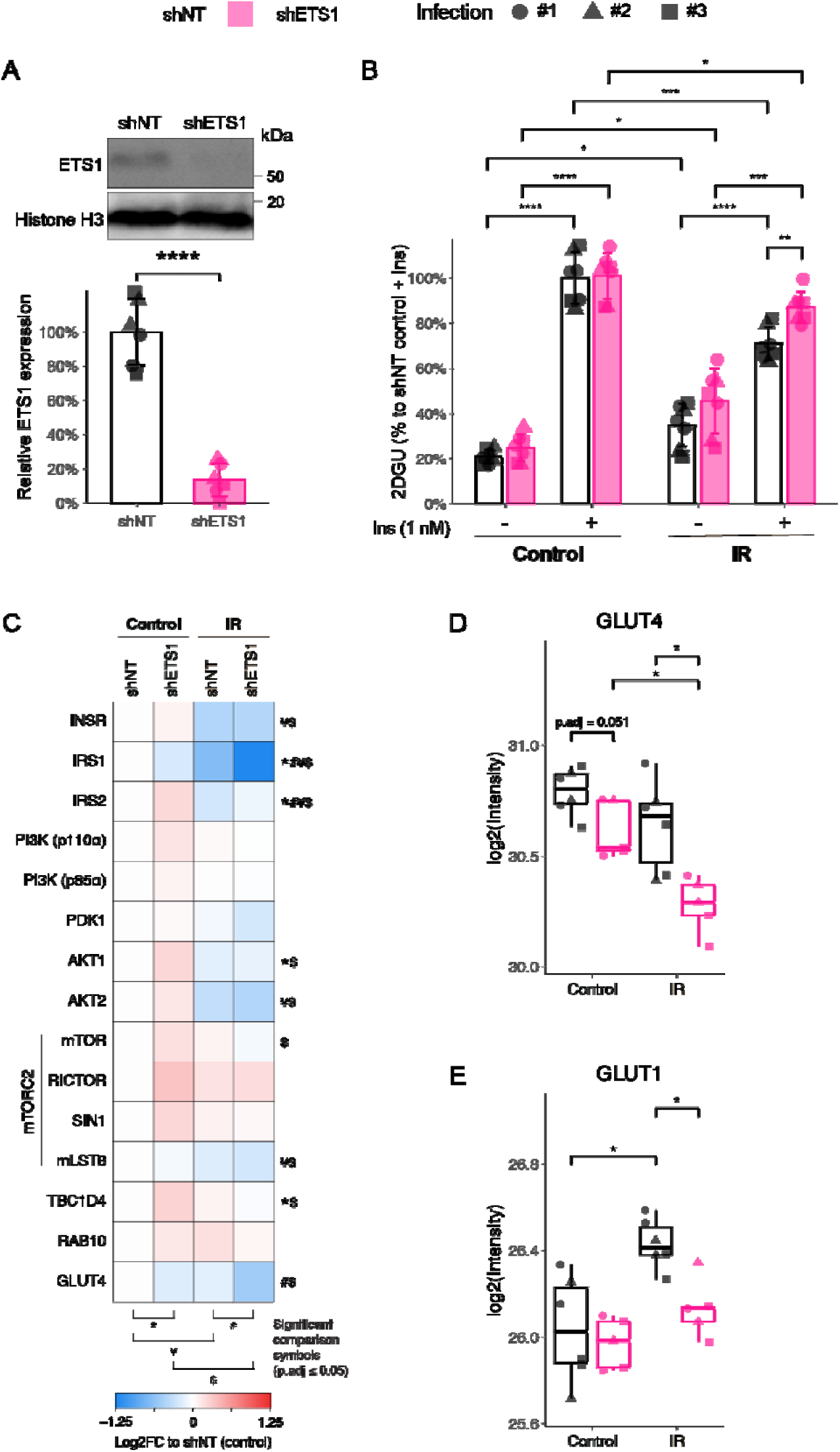
*Ets1* suppression enhances insulin-stimulated glucose uptake in insulin-resistant 3T3-L1 adipocytes. **(A)** ETS1 protein levels in 3T3-L1 adipocytes expressing short hairpin RNA targeting *Ets1* (shETS1) or a non-targeting control (shNT) across multiple independent retroviral infections. The representative immunoblot (top) and quantification (bottom) are shown. Data are mean ± SD; a t-test was used to assess the mean differences between groups (****p ≤ 0.0001). Nuclear extraction was performed, and histone H3 was used as the loading control. **(B)** Basal and insulin-stimulated 2DGU (at 1 nM) in shNT and shETS1 cells under control or insulin-resistant (IR) conditions. Data are mean ± SD; asterisks denote the statistical significance of mean differences between groups (t-test; Benjamini–Hochberg correction; *p.adj ≤ 0.05, **p.adj ≤ 0.01, ***p.adj ≤ 0.001, ****p.adj ≤ 0.0001). **(C)** Canonical proteins that regulate insulin-stimulated glucose uptake. Various symbols were used to denote the statistical significance of mean differences between groups as indicated (t-test; Benjamini-Hochberg adjustment within protein; *^#^¥^$^ p.adj ≤ 0.05). **(D-E)** Protein expression of GLUT4 (D) and GLUT1 (E). Data are presented as box plots. Asterisks denote the statistical significance of mean differences between groups (t-test, Benjamini-Hochberg adjustment; *p.adj ≤ 0.05).

ETS1 KD had no detectable effect on adipocyte differentiation (Extended Data Fig. 4A), nor on basal or insulin-stimulated 2DGU in control adipocytes (Extended Data Fig. 4B). Because the chromosome 9 locus influenced adipose tissue insulin action only in WD-fed mice, we asked whether ETS1 similarly functions only during metabolic stress. Insulin resistance, induced by chronic insulin exposure^38^, reduced insulin-stimulated 2DGU by approximately 30% compared to untreated shNT controls (Fig. 5B). Strikingly, *Ets1* silencing restored 55% of this defect (Fig. 5B).

To investigate the underlying mechanisms, we performed deep proteomic profiling of control and insulin-resistant adipocytes with or without *Ets1* silencing (Extended Data Fig. 4C). Principal component analysis (PCA) revealed a pronounced proteomic shift in insulin-resistant shETS1 samples relative to all other groups (Extended Data Fig. 4D), indicating that ETS1 exerts broad regulatory effects specifically under conditions of metabolic stress, such as insulin resistance.

Importantly, the ETS1 KD-mediated restoration of insulin-stimulated 2DGU could not be explained by changes in canonical insulin signalling pathways (Fig. 5C). Upon insulin resistance, the protein abundance of INSR, IRS1, IRS2, and AKT2 was similarly decreased in both shNT and shETS1 cells (Fig. 5C), and phosphorylation of key insulin-signalling proteins remained comparably attenuated (Extended Data Fig. 4E). Similarly, the restored 2DGU with *Ets1* silencing was not due to increased glucose transporter abundance as both GLUT4 and GLUT1 showed a small but significant decrease in insulin-resistant shETS1 cells (Fig. 5D, E). Collectively, these findings establish ETS1 as a context-dependent regulator of adipocyte insulin responsiveness whose functional behaviour closely mirrors the diet-dependent genetic effects observed in *ex vivo* WAT.

### ETS1 remodels heme metabolism during metabolic stress

To identify the mechanisms by which ETS1 regulates adipocyte insulin action, we performed differential expression analysis across all experimental conditions (Fig. 6A, B). Proteomic remodelling differed markedly between genotypes. Whereas insulin resistance altered only 219 proteins (2.8%) in shNT adipocytes, ETS1 depletion remodelled 3,372 proteins (43.7%) under the same conditions, again demonstrating ETS1’s stress-responsive role.

**Figure 6.**
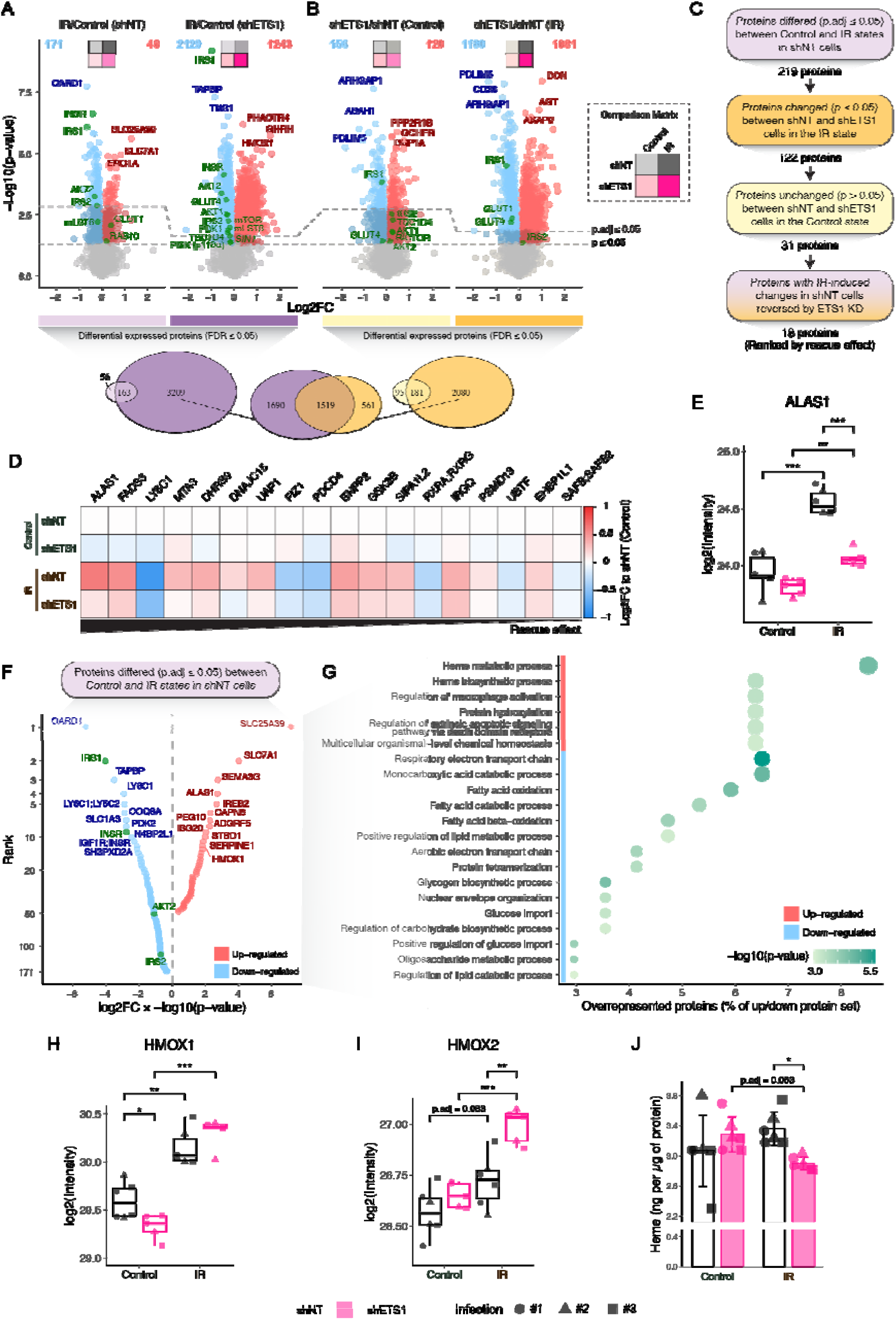
Proteomic analysis reveals *Ets1*-dependent regulation of heme metabolism under insulin resistance. **(A-B)** Multiple t-tests were performed between different experimental groups for each protein. The comparison matrix illustrates the comparisons performed. Dashed lines indicate significance thresholds (p ≤ 0.05, Benjamini-Hochberg adjustment: p.adj ≤ 0.05). For each comparison, the number of differentially expressed proteins (p.adj ≤ 0.05) is printed. Selected top hits, along with GLUT1 and canonical GLUT4 trafficking proteins (p ≤ 0.05), are annotated. Venn diagrams (bottom) show overlap between significant protein sets. **(C-D)** Workflow to identify proteins that changed significantly with insulin resistance (IR) in shNT cells and subsequently returned towards their control levels by silencing *Ets1*. Proteins are ranked by the magnitude of rescue (D). **(E)** ALAS1 protein abundance in shNT and shETS1 cells under control or IR conditions. **(F-G)** Proteins altered by IR in shNT cells (F), and over-representation analysis performed on these proteins using Gene Ontology Biological Process terms (G). The x-axis and the size of the circle denote the percentage of input proteins annotated to a pathway. **(H-J)** Protein expression of HMOX1 (H), HMOX2 (I), and the total heme level measured in shNT and shETS1 cells under control or IR conditions (J). For (E, H-J), asterisks denote the statistical significance of mean differences between groups (t-test, Benjamini-Hochberg adjustment; *p.adj ≤ 0.05, **p.adj ≤ 0.01, ***p.adj ≤ 0.001). Data in (E, H-I) are presented as box plots, while (J) shows mean ± SD.

To identify potential mediators of the rescue phenotype, we focused on proteins that were significantly changed by insulin resistance and restored towards control cell levels following ETS1 KD (Fig. 6C). This analysis identified 18 candidate proteins whose expression pattern matched the 2DGU phenotype (Fig. 6D). This included known regulators of glucose uptake, such as GSK3B^39^ and EHBP1L1^40^.

Among these, the strongest candidate was 5’-Aminolevulinate synthase 1 (ALAS1) (Fig. 6D), the rate-limiting enzyme in heme biosynthesis. ALAS1 abundance increased by approximately 50% in insulin-resistant shNT cells but was completely normalised with ETS1 KD (Fig. 6E). Interestingly, heme biosynthetic and metabolic pathways were upregulated upon induction of insulin resistance (Fig. 6F, G).

These observations led us to investigate whether ETS1 also remodels heme turnover. While the heme-degrading enzyme HMOX1 was upregulated in response to insulin resistance independently of ETS1 manipulation (Fig. 6H), HMOX2 was selectively upregulated in insulin-resistant shETS1 cells (Fig. 6I), suggesting enhanced heme degradation. Consistent with these proteomic changes, total cellular heme content was significantly decreased in insulin-resistant shETS1 cells relative to shNT cells (Fig. 6J). Collectively, these findings suggest a coordinated remodelling of heme metabolism by ETS1 during metabolic stress.

### ETS1 coordinates the heme–iron axis during metabolic stress

Because heme synthesis and degradation are intimately coupled to cellular iron homeostasis^41^, and excess labile iron can contribute to oxidative stress and insulin resistance by promoting the generation of reactive oxygen species (ROS) via the Fenton reaction^42^, we next investigated whether ETS1 also remodels cellular iron handling.

Interestingly, the abundance of several proteins involved in iron uptake and storage was changed under insulin-resistant conditions and ETS1 KD (Fig. 7A-D). Most notably, the expression of the transferrin receptor (TfR) and divalent metal transporter 1 (DMT1), two key mediators of cellular iron uptake, exhibited different or even opposite responses to insulin resistance in shNT and shETS1 cells (Fig. 7A, B). While insulin resistance significantly upregulated TfR expression and had no effect on DMT1 in shNT cells, the same insulin-resistant conditions significantly downregulated both TfR and DMT1 expression in shETS1 cells, pointing towards lower iron uptake in ETS1 KD cells.

**Figure 7.**
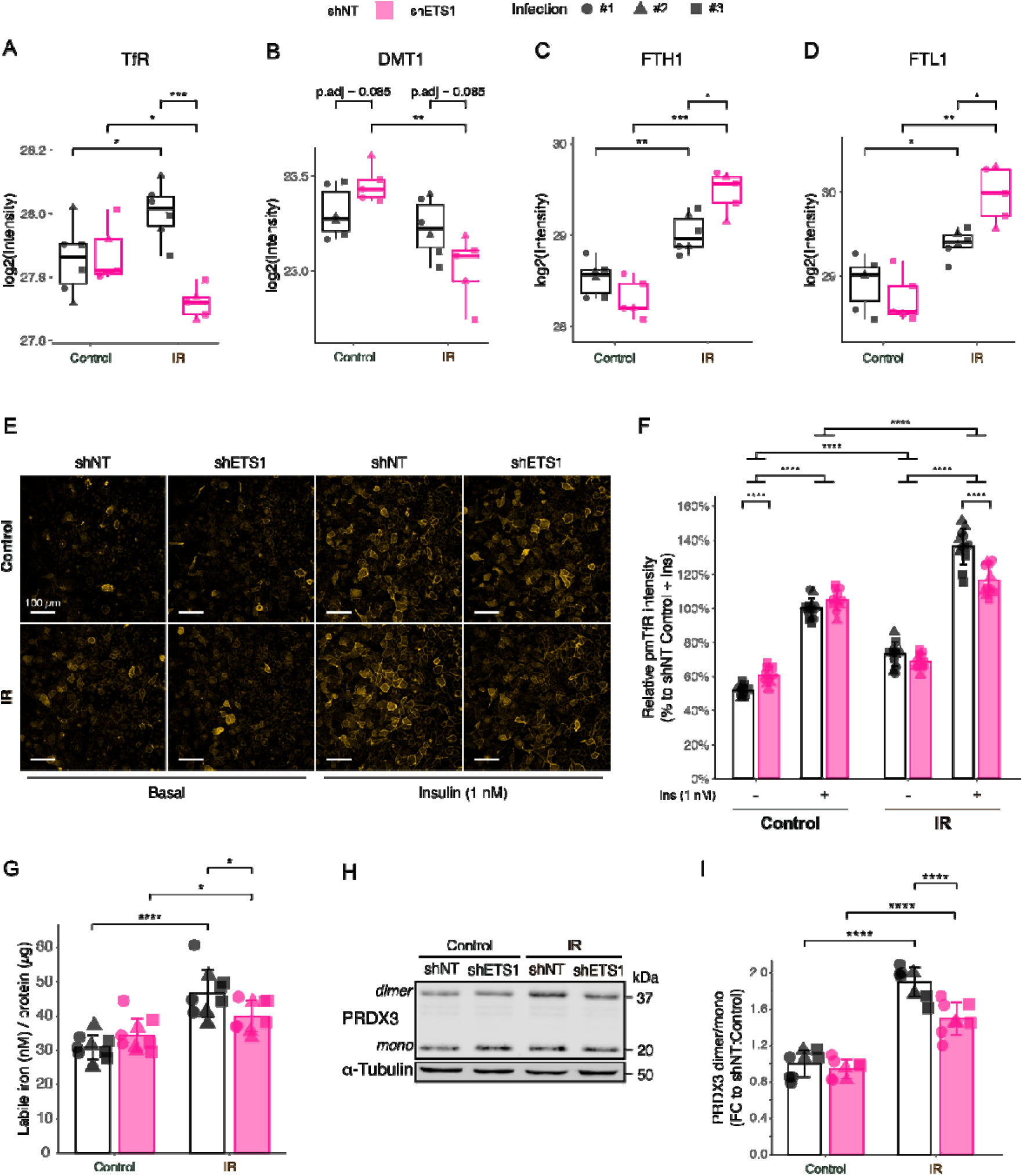
*Ets1* regulates adipocyte iron handling and mitochondrial redox state in a context-dependent manner. **(A-D)** Protein expression of TfR (A), DMT1 (also known as SLC11A2) (B), FTH1 (C) and FTL1 (D) in shNT and shETS1 cells under control or insulin-resistant (IR) conditions. **(E)** Plasma membrane labelling of TfR (pmTfR) in shNT and shETS1 cells under control or IR conditions, treated with and without 1 nM insulin for 20 mins. Scale bar is 100 µm. **(F)** Quantification of pmTfR in (E). **(G)** Labile iron level measured in shNT and shETS1 cells under control or IR conditions. **(H-I)** Mitochondrial oxidative stress in shNT and shETS1 cells under control or IR conditions was assessed by immunoblotting using the dimeric state of PRDX3 as a surrogate marker. The representative blot (H) and quantifications of dimer over monomer ratio (I) are shown. Data in (A-D) are presented as box plots, while (F, G, I) show mean ± SD. Asterisks denote the statistical significance of mean differences between groups (t-test, Benjamini-Hochberg adjustment; *p.adj ≤ 0.05, **p.adj ≤ 0.01, ***p.adj ≤ 0.001, ****p.adj ≤ 0.0001).

As TfR undergoes insulin-dependent translocation from intracellular compartments to the cell surface^43,44^, thereby promoting transferrin-bound iron uptake, we examined insulin-dependent TfR translocation to the plasma membrane (pmTfR) in ETS1 KD cells. As expected^38^, acute insulin stimulation increased pmTfR levels in shNT adipocytes (Fig. 7E, F). Consistent with the upregulation of TfR, insulin resistance potentiated rather than impaired this response, resulting in a further increase in insulin-stimulated pmTfR levels in shNT cells. *Ets1* silencing significantly attenuated this effect, restoring pmTfR levels towards those observed in control cells (Fig. 7E, F). These findings suggest that insulin resistance increases insulin-stimulated iron uptake and that ETS1 is involved in this adaptation.

In addition to controlling iron uptake, cells protect against iron-induced toxicity by sequestering excess iron within ferritin complexes^41,45^. Accordingly, expression levels of both ferritin heavy chain (FTH1) and ferritin light chain (FTL1) were significantly increased under insulin-resistant conditions and further elevated following ETS KD (Fig. 7C, D), suggesting enhanced iron storage. Consistent with the combined effects on iron uptake and storage, labile iron levels were increased in insulin-resistant shNT cells but restored towards control levels following *Ets1* silencing (Fig. 7G).

Given that labile iron can promote reactive oxygen species (ROS) production, we next assessed a readout of oxidative stress. Insulin resistance increased mitochondrial ROS, measured by dimerisation of the mitochondrial peroxiredoxin PRDX3 (Fig. 7H, I), as previously shown^46–48^, whereas this increase was significantly attenuated in shETS1 cells, coinciding with the restoration of insulin-stimulated 2DGU (Fig. 5B).

To determine whether these effects could result from direct transcriptional regulation via ETS1, we analysed published ETS1 chromatin immunoprecipitation sequencing (ChIP-seq) datasets from adipocytes exposed to cold stress^49^. ETS1 binding was detected at regulatory regions of multiple genes involved in heme-iron metabolism (Extended Data Fig. 5), suggesting that ETS1 acts as a stress-dependent transcriptional activator of *Alas1*, *Tfrc*, and *Dmt1*, while repressing *Hmox2*, *Fth1*, and *Ftl1*. This is consistent with previous reports that ETS1 can switch between transcriptional activator and repressor in a context-dependent manner^50^. These observations support a direct role for ETS1 in coordinating the heme and iron pathways under conditions of metabolic stress.

Together with the genetic data, these findings establish the heme–iron axis as the principal downstream effector through which ETS1 links metabolic stress to impaired adipocyte insulin responsiveness. More broadly, they illustrate how direct tissue-level phenotyping coupled with systems genetics can uncover unexpected molecular pathways regulating tissue-specific insulin action.

## Discussion

Metabolic disease arises through complex interactions between genetic variation and environmental exposure^2^, yet identifying the molecular pathways that determine tissue-specific insulin responsiveness has remained a major challenge. Here, we addressed this problem by combining systems genetics with direct measures of adipose tissue insulin action in a genetically diverse mouse population. By quantifying insulin-stimulated glucose uptake directly in adipose tissue, we were able to dissect genetic variation in adipocyte insulin responsiveness independent of adiposity, identifying 39 genetic loci associated with this trait. Functional studies established ETS1 as a regulator of adipocyte insulin action, uncovering a previously unrecognised role for the heme–iron axis in linking metabolic stress to adipocyte insulin resistance. More broadly, these findings demonstrate that deep tissue-specific physiological phenotyping substantially enhances the power of systems genetics to identify causal regulators of complex metabolic traits.

A major conceptual advance emerging from this study is that adipose tissue insulin action exhibits substantial genetic variation that is not explained simply by differences in adiposity. Obesity is widely regarded as the principal determinant of adipose tissue insulin resistance^19,51^, and indeed, we observed a significant inverse relationship between adiposity and insulin-stimulated glucose uptake. However, considerable heterogeneity remained among animals with comparable degrees of adiposity, indicating that obesity alone cannot account for variation in adipose tissue insulin action. By accounting for adiposity during genetic mapping, we were able to distinguish loci associated with adipose tissue insulin action from those influencing obesity itself. This distinction is critical because most human genetic studies necessarily rely on systemic traits, such as fasting insulin^10,11^, body fat distribution^52,53^ or BMI^54^, which represent integrated outputs of multiple organs and physiological processes. Our findings therefore highlight a broader principle for systems genetics: increasing the physiological resolution of complex traits can markedly improve the ability to identify causal genes and pathways. Similar approaches may prove valuable for dissecting tissue-specific insulin resistance in skeletal muscle, liver and other metabolically important organs.

Among the loci identified, the chromosome 9 locus containing *Ets1* was particularly compelling because the effects of the lead SNP were independent of adiposity yet manifested only under WD feeding. Functional studies closely recapitulated these genetic observations. *Ets1* silencing in 3T3-L1 cells had no effect on glucose uptake in control adipocytes but restored insulin-stimulated glucose uptake under insulin-resistant conditions, demonstrating that ETS1 functions as a context-dependent regulator of adipocyte insulin responsiveness. Previous studies have implicated ETS1 in immune activation, vascular biology, cancer development, and diverse metabolic functions^55–63^. In adipose tissue, Wu et al. demonstrated that ETS1 expression in isolated adipocytes increases during high-fat feeding and that adipocyte-specific deletion of ETS1 protects mice from diet-induced metabolic dysfunction^49^. Our findings substantially extend this work by identifying the molecular pathway linking ETS1 to adipocyte insulin action, establishing ETS1 as a key mediator of the adipocyte response to metabolic stress.

A second major mechanistic insight emerging from this study is the identification of the heme-iron axis as a downstream effector of ETS1 during insulin resistance. Proteomic profiling revealed coordinated remodelling of enzymes that control heme biosynthesis and degradation, together with proteins controlling iron uptake and storage. *Ets1* silencing reduced the abundance of proteins involved in iron acquisition while simultaneously increasing ferritin subunits involved in safe iron sequestration. Importantly, ETS1 ChIP-seq data revealed stress-induced ETS1 occupancy at regulatory regions of key genes within these pathways, supporting a direct role for ETS1 in their gene expression regulation. At the biochemical level, *Ets1* silencing reduced cellular heme content, labile iron levels and mitochondrial oxidative stress. Collectively, these findings support a model in which ETS1 orchestrates a stress-responsive transcriptional programme that remodels the heme–iron axis, promoting cellular oxidative stress and impaired adipocyte insulin action during metabolic stress.

These findings substantially extend a growing body of evidence linking dysregulated iron metabolism to insulin resistance and metabolic disease^64–68^. Iron overload disorders, such as hemochromatosis, are frequently associated with insulin resistance and impaired glucose homeostasis^69^, while therapeutic phlebotomy improves insulin sensitivity and glucose tolerance in individuals with elevated ferritin levels^67,70,71^. Gabrielsen and colleagues further demonstrated that adipocyte iron directly regulates systemic insulin sensitivity, showing that increased adipocyte iron suppresses adiponectin expression, with phlebotomy simultaneously increasing circulating adiponectin and improving glucose tolerance in humans^72^. Subsequent human studies reported that circulating ferritin and other markers of iron levels correlate with adipocyte insulin resistance independently of obesity and systemic inflammation^73^. More recently, attention has shifted from total cellular iron to the labile iron pool, which promotes the generation of reactive oxygen species through Fenton chemistry, driving oxidative stress, mitochondrial dysfunction and lipid peroxidation^42^. Our findings extend this framework by identifying ETS1 as an upstream transcriptional regulator of adipocyte iron homeostasis. Rather than viewing iron accumulation as a secondary consequence of obesity, our data support a model in which genetically determined remodelling of the heme-iron axis increases intracellular labile iron during metabolic stress, thereby promoting oxidative stress and adipocyte insulin resistance.

Interestingly, ETS1 is a well-characterised downstream target of the RAS–ERK signalling pathway^74,75^, where ERK-mediated phosphorylation enhances its transcriptional activity through recruitment of the co-activators CBP and p300^76^. This raises the possibility that nutrient- or inflammation-induced MAPK signalling activates ETS1 during metabolic stress, thereby coupling extracellular stress signals to remodelling of the heme–iron axis. This is an interesting concept for future study.

Several limitations of this study should be considered. First, adipose tissue insulin action was quantified *ex vivo* and therefore does not capture all physiological influences present *in vivo.* However, this approach also represents a major strength because it enabled precise quantification of insulin dose-response independent of confounding variables such as tissue perfusion and blood flow^21^, thereby greatly improving the sensitivity of genetic mapping.

Second, mechanistic studies were performed in 3T3-L1 adipocytes, which do not fully recapitulate the complexity of adipose tissue *in vivo*. Nevertheless, previous results from adipocyte-specific *Ets1* knockout mice^49^ are highly consistent with our findings. Finally, although our data strongly implicate the heme-iron axis as a downstream mediator of ETS1 action, direct manipulation of these pathways will be required to establish definitive causality. Nonetheless, heme administration during human adipocyte differentiation has been reported to increase the expression of iron-overload markers and impair insulin-stimulated glucose uptake^77^, providing independent support for our proposed model.

In conclusion, this study establishes deep tissue-specific physiological phenotyping combined with systems genetics as a powerful strategy for identifying regulators of tissue-specific insulin action. By integrating physiological phenotyping, genetic mapping, and quantitative proteomics, we identify ETS1 as a diet-dependent regulator of adipocyte insulin action, acting through the heme–iron axis. More broadly, these findings demonstrate that increasing the physiological resolution of complex metabolic traits substantially enhances the power of systems genetics to identify causal pathways that remain hidden when relying solely on conventional whole-body phenotypes. This framework provides a general strategy for dissecting the genetic architecture of tissue-specific metabolism and should facilitate the discovery of novel therapeutic targets across a broad range of metabolic diseases.

## Methods

### Mouse housing, feeding and phenotyping

All animal experiments were conducted following NHMRC (Australia) guidelines for animal research and were approved by The University of Sydney Animal Ethics Committee. All mice were group-housed at 23°C on a 12-h light/dark cycle (0600-1800) with free access to water and food.

#### Inbred animals

C57BL/6J (BL6), DBA/2J (DBA), A/J, and NOD/ShiLtJ (NOD) inbred mouse strains were obtained from the Ozgene Animal Resources Centre (Perth, WA, Australia); 129X1/SvJ (129), CAST/EiJ (CAST), and WSB/EiJ (WSB) were obtained from Australian BioResources (Moss Vale, NSW, Australia). Mice were provided ad libitum access to a standard laboratory chow diet (Gordon’s Specialty Stock Feeds, NSW, Australia) and were used for experiments at 16-18 weeks of age.

#### Diversity Outbred in Australia (DOz)

DOz mice were bred and housed at the Charles Perkins Centre, University of Sydney, NSW, Australia, as previously described^20,78,79^. The DOz mice used in this study were outbred for 27 to 35 generations and included a total of 559 animals across 9 cohorts (435 males, 124 females). Genomic DNA was extracted from each mouse and analysed using SNP genotyping with the Giga Mouse Universal Genotyping Array from Neogen (Ipswich, QLD, Australia), followed by diagnostics and cleaning as described^80^. Starting at 10 weeks of age, mice were given *ad libitum* access to either chow (n = 277; 242 males, 35 females) or a Western diet (WD) for 8 weeks (WD8; n = 80; male only) or 14 weeks (WD14; n = 202; 113 males, 89 females). The WD was prepared in-house and contained approximately 61% of calories from fat, 19% from carbohydrate, and 20% from protein, with the following composition (% w/w): 25.5% casein, 0.4% L-cystine, 16.0% cornstarch, 9.3% sucrose, 6.4% α-cellulose, 31.3% lard, 3.2% safflower oil, 1.0% cholesterol, 6.4% AIN-93 mineral mix (MP Biomedicals), 1.3% AIN-93 vitamin mix (MP Biomedicals), and 0.3% choline.

#### Systemic metabolic measures in the DOz

Cull weight was measured at the end of the diet intervention upon animal sacrifice. Fat and lean mass were measured using EchoMRI-900 (EchoMRI Corporation Pte Ltd, Singapore) at the age of 14 weeks for the chow and WD8 groups, and at 23 weeks for the WD14 group. Glucose tolerance was assessed through an oral glucose tolerance test (GTT) in fasting mice for 6 hours prior to administering a 20% glucose solution in water at 2 mg/kg of lean mass. Oral GTT was performed at the age of 14 weeks for the chow and WD8 groups, and at 23 weeks for the WD14 group. Blood glucose levels were taken directly from tail blood using a handheld glucometer (Accu-Chek, Roche Diabetes Care, NSW, Australia) at 0, 15, 30, 45, 60, and 90 min after glucose gavage. Blood insulin levels at 0 and 15 min were measured using mouse insulin ELISA kits from Crystal Chem USA (Elk Grove Village, IL, USA), following the manufacturer’s instructions. Triglycerides were extracted from liver tissue by adding 800 µL of chloroform and methanol (2:1 ratio), vigorously mixing on a rocking platform for 30 min, then adding 400 µL of 0.6% NaCl. Samples were centrifuged at 3000 rpm for 15 min, and the bottom phase was collected. The chloroform-methanol extraction was repeated, and the bottom phase was added to the same tube as the first extraction. Samples were dried down in a Gene-Vac (HPLC mode at 37°C) for 2 hours, then resuspended in isopropanol and warmed to 55°C for use in a colourimetric triglyceride assay (Thermo Fisher Scientific, Cat. #981786).

#### The high-throughput *ex vivo* 2DGU assay

Basal and insulin-stimulated glucose uptake in gWAT was assessed using radiolabelled 2-Deoxy-D-glucose (2DG), in which 2DG uptake (2DGU) serves as a proxy for glucose uptake. The detailed procedure of this assay is illustrated in Supp. Fig. 1.

Basal media containing Gibco^TM^ DMEM powder, high glucose (Thermo Fisher Scientific, Cat. #12800017), 2% BSA (Bovogen, Cat. #BSAS 1.0) and 25 mM HEPES (Sigma-Aldrich, Cat. #H4034) at pH 7.4 was prewarmed to 37°C. gWAT from each individual mouse was excised and placed into a 23 mL flat-bottom tube with a cap (Sarstedt, Cat. #58.490 and #65.790) containing basal media. Next, gWAT explants were chopped into uniform fine pieces (∼2-3 mm) using iris dissection scissors (World Precision Instruments, Cat. #503261) in the basal media. Then, the explants were washed three times with warmed basal media by allowing them to float between media additions, and the media below was removed with an 18G blunt draw-up needle (Terumo, Cat. #219894) and a 10 mL syringe. The explants were placed in the tube containing 10 mL of basal media on a rocking platform in a 37°C water bath and gently shaken. The interval between the first and last samples was kept to a maximum of 2 hours. After completing the above steps on the last animal, all explants were left incubated for 2 hours.

After the 2-hour incubation in the basal media, explants were washed twice with 10 mL of room-temperature (RT) PBS (Thermo Fisher Scientific, Cat. #18912014) to remove the glucose-containing basal media, using the same float-and-18G-blunt-needle technique described above. They were then washed with 10 mL of pre-warmed assay buffer containing 120 mM NaCl, 600 µM Na_2_HPO_4_, 6 mM KCl, 400 µM NaH_2_PO_4_, 1.2 mM MgSO_4_, 25 mM HEPES, 1 mM CaCl_2_, and 2% BSA at pH 7.4. Excess assay buffer was removed, and 50 μL of explants were aliquoted, using a cut 200 μL tip, into a 96-well MultiScreen® High Volume Filter Plate (Merck Millipore, Cat. #MVFCN1225) containing pre-warmed assay buffer. An adhesive film was placed at the bottom of the plate prior to explant and buffer addition to prevent flow-through during the assay. Insulin was added to the wells using an 8-channel pipette at the desired concentrations (0.01 nM [considered as the basal dose], 0.5 nM, 2.5 nM, or 10 nM) for 20 min, and the plate was incubated at 37°C. A mastermix of 2DG was added to each well for the last 5 min of insulin stimulation, resulting in a final mix of 50 µM 2DG (Merck, Cat. #D6134), 2 µCi/mL [^3^H]2DG (Deoxy-D-glucose, 2-[1,2-^3^H(N)]; PerkinElmer, Cat. #NET328001MC), 0.28 µCi/mL [^14^C]Mannitol (Mannitol, D-[2-^14^C]; PerkinElmer, Cat. #NEC852050UC). The total volume per well was 250 µL.

After 20 min of stimulation, 1 mL of ice-cold PBS was added to each well using an 8-channel pipette, and the plate was placed on ice. The adhesive seal was removed from the base of the filter plate, and the plate was placed on a MultiScreen® High Volume Collection Plate (Merck Millipore, Cat. #MVCPN0025). This plate assembly was centrifuged at 100 × g for 2 min, and then the explants were washed a further three times with 1 mL of ice-cold PBS, increasing the centrifugation time by 1 minute and the speed by 100 × g for each subsequent wash step, and emptying the collection plate each time. The adhesive film was placed back at the bottom of the filter plate containing explants, and 250 μL of 100 mM NaOH was added to each well. The explants were incubated at 95°C for 30 min to lyse. Then, the adhesive film was gently removed again, and the filter plate was placed on a new collection plate, which was then centrifuged at 400 × g for 2 min to collect the lysate. The explant lysates were analysed using a BCA assay (Thermo Fisher Scientific, Cat. #23225) for protein normalisation and measured using a Tri-Carb 2810R Liquid Scintillation Counter (PerkinElmer) for ^3^H and ^14^C radiation counts. A fixed volume of the 2DG mastermix was also counted on the scintillation counter to determine ^3^H and ^14^C DPM (disintegrations per minute) per unit volume or per nmol of 2DG.

To measure each explant’s 2DGU, the extracellular volume (ECV) of each lysate sample was first calculated using the sample’s ^14^C DPM counts and the corresponding ^14^C DPM-per-volume conversion factor obtained from the 2DG mastermix. Extracellular 2DG (EC-2DG; expressed as ^3^H DPM) for each sample was then calculated using the ^3^H DPM-per-volume conversion factor and the ECV determined. Intracellular 2DG (IC-2DG; expressed as ^3^H DPM) was next determined by subtracting EC-2DG from the total ^3^H counts. IC-2DG values were subsequently converted from ^3^H DPM to nmol of 2DG using the corresponding ^3^H DPM-per-nmol conversion factor determined from the 2DG mastermix. These IC-2DG were further normalised to the protein amount in each sample and the 2DG uptake time (5 min). The resulting values were considered as the 2DGU measures. For each sample, 2DGU was calculated as follows:

*1)* 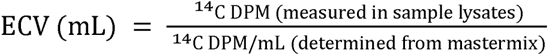
*2)* EC-2DG ^3^H DPM ECV ^3^H DPM/mL determined from mastermix
*3)* IC-2DG ^3^H DPM total ^3^H counts DPM ! EC-2DG ^3^H DPM
*4)* 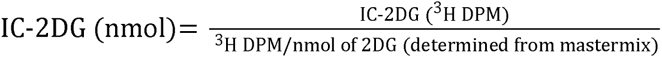
*5)* IC-2DG nmol/mg/hr IC-2DG nmol $ protein amount mg $ incubation time hr

### In vitro work

#### Cell culture, retroviral shRNA transduction, and insulin resistance induction

As previously described^81^, Plat-E cells cultured in 10 cm dishes were transiently transfected with retroviral vector pBABE using Lipofectamine 2000 (Invitrogen, Cat. #52887) according to the manufacturer’s instructions. The medium was replaced the next day with 7 ml per dish. Virus-containing medium was collected after two days, one day at 37°C and another at 32°C, then filtered through a 0.45 µm filter and used immediately for infection or stored at -80°C.

Mycoplasma-free 3T3-L1 fibroblasts were obtained from Howard Green (Harvard Medical School)^82^ and cultured in Dulbecco’s modified Eagle medium (DMEM, high glucose) (Gibco, Cat. #11960-044) supplemented with 10% (v/v) fetal bovine serum (FBS) (Sigma Life Science, Cat. #F9423-500 Ml) and 1× GlutaMAX (Gibco, Cat. #35050-061).

3T3-L1 fibroblasts were infected with puromycin-resistant retroviral vector pBABE encoding shRNA against ETS1 (shETS1; target sequence: 5’ – GCAGACAGACTACTTTGCCAT – 3’)^83^ or non-targeting shRNA (shNT) against luciferase. Three independent viral infections were performed to ensure the reliability and validity of subsequent experiments. For each independent infection, multiple passages were also produced as biological replicates. ETS1 KD efficiency was assessed by immunoblotting after nuclear extraction. Puromycin concentration (2 µg/mL) was maintained during culture of shNT and shETS1 fibroblasts to ensure stable shRNA expression. The fibroblasts were then differentiated into adipocytes upon confluence, as previously described^84^. Brightfield images of shNT and shETS1 adipocytes were acquired using a Millipore DCI Digital Cell Imager. To induce insulin resistance, shNT and shETS1 adipocytes were incubated with a single dose of 10 nM insulin for 24 hrs, as previously described^38,84^.

#### Nuclear extraction

To assess ETS1 knockdown (KD) efficiency, nuclear extraction of shNT and shETS1 cells was performed as previously described^85^, with minor modifications. Briefly, cells washed and pelleted (with ice-cold PBS) were resuspended in 10 mM HEPES (pH 7.9), 10 mM KCl, 0.1 mM EDTA, 0.1 mM EGTA, supplemented with 1 mM DTT and a protease inhibitor cocktail (Roche Applied Science, Cat. #11873580001), and incubated on ice for 15 min to allow swelling. NP-40 (Sigma-Aldrich, Cat. #I3021) was then added (to a final concentration of 0.6%), followed by vigorous vortexing for 10 s. Samples were centrifuged at 13,000 × g for 30 s at 4°C to separate the nuclear and non-nuclear fractions. The nuclear pellet was resuspended in 100 µL 2% SDS in HES buffer (250 mM sucrose, 20 mM HEPES, 1mM EDTA; pH 7.4) containing protease inhibitors, incubated at 60°C for 5 min, and sonicated. Samples were then centrifuged at 13,000 × g for 5 min at 6°C, and the supernatant was collected as the nuclear extract. Protein concentrations were determined using the BCA assay. For immunoblotting, 25 µg of protein (the nuclear fraction) was loaded per sample.

#### Cellular 2DGU assay

shNT and shETS1 adipocytes (under control or insulin-resistant conditions) were serum-starved for 2 h in DMEM/GlutaMAX with 0.2% BSA (pH 7.4) at 37°C, 10% CO_2_. Afterwards, cells were washed three times and incubated for 5 min in prewarmed Krebs-Ringer phosphate (KRP) buffer (0.6 mM Na_2_HPO_4_, 0.4 mM NaH_2_PO_4_, 120 mM NaCl, 6 mM KCl, 1 mM CaCl_2_, 1.2 mM MgSO_4_, and 12.5 mM HEPES; pH 7.4) containing 0.2% BSA. Adipocytes were then stimulated with the desired insulin concentrations for 20 min. To account for nonspecific 2DGU, 50 μM cytochalasin B (in ethanol; Sigma-Aldrich, Cat. #C6762) was added to selected representative wells receiving insulin prior to the addition of 2DG. For the last 5 min of insulin stimulation, 2DG (50 µM 2DG; 0.25 μCi [^3^H]2DG) was added to each well. Cells were then rapidly cooled on ice, washed five times with ice-cold PBS, and permeabilised with 1% Triton X-100 (v/v in PBS, gently shaken for 1 hour). 2DGU was measured via liquid scintillation counting using a TriCarb 2900TR (PerkinElmer). Data were normalised to protein content and reaction time, and further mean-normalised within each experimental plate prior to combining datasets for analysis.

#### Assessment of phosphorylation of key insulin-signalling proteins

shNT and shETS1 adipocytes (under control or insulin-resistant conditions) were serum-starved for 2 h in DMEM/GlutaMAX with 0.2% BSA (pH 7.4), followed by stimulation with or without 1 nM insulin at 37°C, 10% CO_2_. After 10 mins, cells were then rapidly cooled on ice, washed three times with ice-cold PBS, and lysed by sonication in 2% SDS in PBS, containing protease inhibitors and phosphatase inhibitors (2 mM Na_3_VO_4_, 1 mM Na_4_P_2_O_7_, 10 mM NaF), followed by sonication (3 s on/off for 30 s). Samples were centrifuged at 21,000 × g for 15 min at 4°C to remove the floating lipid layer (fat cake). The remaining lysate was then heated at 65°C until clear, followed by a second centrifugation at 21,000 × g for 15 min at 12°C to remove cellular debris. The supernatant was collected, and protein concentrations were determined using the BCA assay. For immunoblotting, 20 µg of protein was loaded per sample.

#### Proteomics sample preparation

shNT and shETS1 adipocytes grown in a 6-well plate (under control or insulin-resistant conditions) were washed three times in ice-cold PBS before scraping into lysis buffer (4% sodium deoxycholate (SDC), 100 mM Tris, pH 8.5) on ice. Lysates were immediately boiled at 95°C for 10 min and sonicated for 1 min using a tip-probe sonicator. Lysates were then centrifuged for 10 min at 20,000 × g at RT, and the supernatant was transferred to 5 mL tubes while avoiding the fat cake. Proteins were purified by high-volume chloroform-methanol precipitation. Briefly, 1600 µL of methanol, 800 µL of chloroform, and 800 µL of water were added sequentially, with brief vortexing between additions. Samples were centrifuged at 2,000 × g for 5 min to achieve phase separation. The aqueous phase was removed, and 2.4 mL of methanol was added to wash the pellet. Samples were centrifuged at 2,000 × g for 5 min to pellet the protein, and the entire supernatant was removed. The protein pellet was resuspended in 300 µL lysis buffer, followed by sonication and centrifugation at 20,000 × g for 10 min at RT to remove insoluble proteins. Protein concentration of the supernatant was determined by the BCA assay. 5 µg of protein was aliquoted into a 1000 µL deep-well plate, and concentrations were adjusted by making up volumes to 40 µL with lysis buffer. Reduction/alkylation buffer (10 mM TCEP, 40 mM 2-chloroacetamide) was added, and the samples were heated for 10 min at 60°C. Samples were then adjusted to 1% SDC through the addition of 120 µL mass spectrometry (MS)-grade H_2_O, and 0.2 mg trypsin and 0.2 mg LysC were added to each sample. Samples were then incubated overnight (18 h) at 37°C.

Peptide purification was performed by StageTip cleanup using SDB-RPS solid-phase extraction material^86^ (Merck; Cat. #66886-U). 160 µL 1% trifluoroacetic acid (TFA) in ethyl acetate was added to samples, and samples were vortexed to stop digestion and to dissolve any precipitated SDC. Samples were centrifuged at 2,000 × g for 1 min to achieve phase separation. The bottom phase was then transferred to the StageTips and loaded by centrifugation at 1,000 × g for 5 min. Stage tips were washed with subsequent spins at 1,000 × g for 5 min with 100 µL 1% TFA in ethyl acetate, 1% TFA in isopropanol, and 0.2% TFA in 5% acetonitrile (ACN). Samples were eluted by the addition of 100 µL 60% ACN with 5% NH_4_OH. Samples were evaporated by vacuum centrifugation, and peptides were reconstituted in 40 µL 5% formic acid.

#### Proteomics analysis

Samples were analysed using a Vanquish Neo UHPLC system coupled to a Q-Exactive HFX mass spectrometer (Thermo Fisher Scientific). 500 ng of peptide sample was injected onto an in-house packed 75 µm x 55 cm column (1.9 µm particle size, ReproSil Pur C18-AQ) and separated using gradient elution, with Buffer A consisting of 0.1% formic acid in water and Buffer B consisting of 0.1% formic acid in 80% ACN. Samples were loaded to the column at a flow rate of 0.4 mL/min at 100% Buffer A for 12 min, before ramping to 40% Buffer B over 68 min, then to 98% Buffer B over 15 min and held for 5 min. Eluting peptides were ionised by electrospray with a spray voltage of 2.4 kV and a transfer capillary temperature of 300°C. Mass spectra were collected using a data-independent acquisition (DIA) method with varying isolation width windows (widths of 27 to 589 m/z) between 350 and 1650 according to Supp. Table 1. MS1 spectra were collected between m/z 350 and 1650 at a resolution of 120,000. Ions were fragmented with a higher-energy collisional dissociation (HCD) collision energy of 25%, and MS2 spectra were collected between m/z 350 and 1650 at a resolution of 30,000, with an automatic gain control (AGC) target of 3e6 and the maximum injection time set to automatic.

#### Total heme and labile iron measurement

##### Adipocytes lysates collection

shNT and shETS1 adipocytes (under control or insulin-resistant conditions) were serum-starved for 2 h in DMEM/GlutaMAX with 0.2% BSA at 37 °C, 10% CO_2_. Cells were then washed three times with ice-cold PBS and lysed on ice in radioimmunoprecipitation assay (RIPA) lysis buffer (50 mM Tris–HCl, 150 mM NaCl, 0.2% SDS (v/w), 0.5% SDC, 1% Triton X-100; pH 7.4) containing protease inhibitors, and homogenised by sonication. After centrifugation at 15,000 × g for 15 min at 6°C, the protein concentration of the clear cell lysate was measured and used for subsequent heme and iron measurements.

##### Fluorescence heme assay

Total heme levels in adipocyte lysates were quantified using a fluorescence-based assay measuring protoporphyrin IX fluorescence following heating in oxalic acid to promote iron release from heme, as previously described^87–89^. Briefly, adipocyte lysates in 10 µL RIPA buffer (40 µg of protein used) were diluted in 100 µL H O and mixed with 1 mL of pre-heated 2 M oxalic acid (Ajax Finechem, Cat. #AJA350). Each sample suspension was split, with half the suspension heated at 100°C for 30 min and the other half kept at RT. Samples were then centrifuged at 21,000 × g for 3 min. Porphyrin fluorescence was measured using a Tecan Infinite® M1000 plate reader (405 nm excitation, 600 nm emission). Total heme content was calculated by subtracting the fluorescence of the unheated suspension from that of the heated suspension. Heme concentrations were determined using a standard curve generated from serial dilutions of hemin chloride (Cayman Chemical, Cat. #16487; prepared by diluting in 0.1 M NaOH) processed identically to the cell samples.

##### Labile iron assay

Labile iron levels in adipocyte lysates were quantified using a colourimetric ferene-based assay measuring the absorbance of the blue Fe² –ferene complex formed in the sample under low ascorbic acid conditions, as previously described^90^. Briefly, adipocyte lysates were prepared in 100 µL RIPA buffer, with 400 µg protein used per sample, followed by the addition of 100 µL 2.5 M ammonium acetate buffer (pH 4.5). The ammonium acetate buffer was prepared from ammonium acetate (Merck, Cat. #A7330) and adjusted to pH 4.5 using glacial acetic acid (Merck, Cat. #695092). Subsequently, 12 µL of iron assay buffer (5 mM ferene (Merck, Cat. #P4272), 10 mM ascorbic acid (Sigma-Aldrich, Cat. #A92902) in ammonium acetate buffer) was added to each sample. Samples were vortexed and incubated overnight at RT, followed by centrifugation at 15,000 × g for 5 min. Supernatants (200 µL) were transferred to a 96-well plate, and absorbance at 595 nm was measured using a Tecan Infinite® M1000 plate reader. Labile iron concentrations were calculated from a standard curve generated using serial dilutions of a 20 mM iron stock solution processed identically to the cell lysates. The iron stock solution was prepared by dissolving carbonyl iron powder (Merck, Cat. #44890) in 37% HCl (Ajax Finechem, Cat. #AJA1367) overnight and diluting to the desired concentration with deionised water, as previously described^91^.

#### Immunofluorescence staining of endogenous TfR translocation

Transferrin receptor (TfR) surface trafficking in shNT and shETS1 adipocytes (under control or insulin-resistant conditions) was assessed by immunofluorescence staining using methods adapted from our previous work^38,84^. Briefly, adipocytes were serum-starved for 2 h in DMEM/GlutaMAX containing 0.2% BSA (pH 7.4), followed by stimulation with or without 1 nM insulin at 37°C for 20 mins. Cells were then washed three times with ice-cold PBS supplemented with 1 mM CaCl and 1 mM MgCl (PBS+/+). Plasma membrane TfR (pmTfR) was labelled by incubating cells for 2 hours at 4°C with a surface antibody solution containing 500 ng/mL anti-TfR (CD71; Rat) (Thermo Fisher Scientific, Cat. #14-0711-82) in 2% horse serum in PBS+/+. Cells were then washed twice with ice-cold PBS+/+, incubated with WGA–FITC lectin (4 µg/mL; Merck, Cat. #L4895) for 20 min at 4°C with foil, and again washed three times with ice-cold PBS+/+. Subsequently, cells were fixed with 4% paraformaldehyde (ProSciTech, Cat. #C004) for 5 min on ice, followed by 15 min at RT, quenched with 50 mM glycine (in PBS+/+; 5 min), and washed with PBS+/+. Secondary antibody solution (2.5 μg/mL anti-Rat Alexa 555 (Invitrogen, Cat. #A21434), Hoechst 33342 (1:10,000; Thermo Fisher Scientific, Cat. #H3570) and 2% horse serum in PBS+/+) were applied for 60 min at RT. Plates were then washed three times with degassed PBS+/+ and prepared in degassed imaging buffer (5% (v/v) glycerol, 2.5% (w/v) 1,4-diazabicyclo[2.2.2]octane; pH 8.5) for imaging. Plates were imaged, and surface TfR levels were quantified using systems described previously^38^, using Harmony (Revvity, Part #HH17000019). Lectin-defined plasma membrane regions were identified by image filtering and masking, after which mean TfR fluorescence intensity within the membrane rim regions was quantified. Data were further mean-normalised within each experimental plate for downstream analysis.

#### Assessment of PRDX3 dimerisation

Mitochondrial ROS in shNT and shETS1 adipocytes was measured by PRDX3 dimerisation, as previously described^47,48^. Briefly, cells were washed three times with ice-cold PBS containing 10 μg/mL catalase and incubated with 100 nM N-Ethylmaleimide (NEM; Tokyo Chemical Industry, Cat. #E0136) in PBS on ice for 10 min. Cells were lysed by sonication in 2% SDS in PBS, containing protease inhibitors and 100 nM NEM. Protein concentration was measured using the BCA assay. 20 µg of protein was used for each sample loading.

#### Immunoblotting

Samples were prepared in a sample buffer containing 1x Laemmli buffer (2% (w/v) SDS,

62.5 mM Tris–HCl (pH 6.8), 10% (v/v) glycerol, 0.1% (w/v) bromophenol blue) supplemented with 50 mM TCEP (Thermo Fisher Scientific, Cat. #77720). Prepared samples were subjected to SDS-PAGE, and the gels were transferred to polyvinylidene difluoride (PVDF) membranes. Membranes were subjected to immunoblotting with primary antibodies as indicated in Supp. Table 2. For detecting ETS1 and histone H3 protein levels, horseradish peroxidase (HRP)-labelled secondary antibodies, followed by detection using enhanced chemiluminescence (ECL) substrates (Thermo Fisher Scientific, Cat. #34577) on the ChemiDoc™ MP Imaging System (Bio-Rad). For other experiments, either infrared dye 700- or 800-conjugated secondary antibodies (Thermo Fisher Scientific, Cat. #A32735 or A21036) were used. Detection was carried out using an Odyssey CLx Imaging System (LI-COR Biosciences). All densitometry analyses were performed using Image Studio 6.0 (LI-COR Biotech). ETS1 band intensities were normalised to the loading control (histone H3), and PRDX3 dimerisation was quantified by calculating the ratio of dimer-to-monomer band intensities. Quantifications were then mean-normalised within each blot prior to combining datasets for analysis.

### Data processing and statistical analyses

All statistical analyses were performed in RStudio using R 4.3.0. Statistical tests applied are specified in the corresponding figure legends. Unless otherwise stated, multiple testing correction was performed with the Benjamini–Hochberg method^92^. Where statistical analyses assumed normality, Box–Cox^93^ or log transformation was applied. Data visualisation was performed using the R/ggplot2 package and other associated R packages.

#### Animal phenotypic analyses

A linear mixed-effects model was used to assess how 2DGU measures were influenced by mouse strain (BL6 and DBA), insulin dose, and experimental day, while accounting for repeated measurements from the same animal. The GTT area of the curve (AOC) was calculated relative to the baseline glucose measure across all time points. Homeostatic Model Assessment for Insulin Resistance (HOMA-IR)^94^ and Matsuda Index^95^ were calculated using standard published equations based on glucose and insulin measurements during GTT^19^. For 2DGU phenotypes, the area under the curve (AUC) was calculated using both basal and insulin-stimulated uptake values, and AOC were calculated using the lowest 2DGU value as the baseline for each animal. Clustering of the 2DGU traits was performed using hierarchical clustering to group animals based on both absolute 2DGU values and the derived pairwise response differences between doses, followed by PCA. Pearson correlation was used to assess the relationships among 2DGU AOC, adiposity, GTT AOC, Matsuda Index, and liver triglyceride content, using Box–Cox transformed data where necessary. Adiposity-adjusted 2DGU AOC was calculated by using the residuals from a linear regression model with 2DGU AOC as the dependent variable and adiposity as the independent variable. A linear model was fitted to assess how 2DGU AOC associates with GTT AOC, Matsuda Index, and liver triglyceride content, with or without adjusting for adiposity.

#### Genetic analysis

Genetic mapping analyses were performed in R using the R/qtl2 package^31^. Raw 2DGU values were Box–Cox transformed prior to mapping^93^. Sex, diet, and adiposity were included as additive covariates when mapping. Kinship matrices were calculated using the leave-one-chromosome-out (LOCO) method. Heritability estimates were calculated using a linear mixed model that accounted for animal kinship, using the qtl2:: est_herit() function. Haplotype-based analysis was performed using founder allele probabilities derived from DOz genotype data, while SNP-based analysis used genotype probabilities at each SNP marker. Genome-wide significance thresholds were determined using 1,000 permutation tests with significance set at p < 0.05, while chromosome-wide significance thresholds were set at p <

0.63. The SNP that was genotyped (on_map = TRUE) with the highest LOD score (lead SNP) at the chromosome 9 locus was further analysed to assess its relationship with 2DGU phenotypes, adiposity, GTT AOC, Matsuda Index, and liver triglyceride content. Linkage disequilibrium (LD) analysis was performed using the R/snpStats package^96,97^. *Synteny*^37^ was used to retrieve human genetic evidence for glycemic traits for the gene of interest, filtered to traits with at least one HuGE score ≥3 across the selected genes.

#### Proteomics data processing and analysis

Proteomics raw data files were searched using DIA-NN 1.9.2 using a library-free FASTA search against the reviewed UniProt mouse proteome (downloaded April 2025) with deep learning enabled^98^. The protease was set to Trypsin/P with 1 missed cleavage, N-term Methionine excision, carbamidomethylation and Methionine oxidation options included. Peptide length was set to 7-30, precursor range 350-1650, and fragment range 350-1650, and FDR set to 1%. Proteomic data were filtered for protein-level missingness, retaining proteins that were reliably quantified in at least one experimental group (with < 50% missingness). Protein intensity values were then log2-transformed and median-normalised by shifting each sample to the global median intensity across all samples.

PCA was performed to assess sample and condition clustering. Differential expression analyses between experimental groups were performed using multiple t-tests, and proteins were classified based on significance thresholds of nominal p ≤ 0.05 and adjusted p (p.adj) ≤ 0.05 (Benjamini–Hochberg method) for each comparison. Protein changes between conditions were visualised using heatmaps, volcano plots, and box plots where indicated. Pathway over-representation analysis was performed using the R/clusterProfiler package^99^ with Gene Ontology (GO) Biological Process terms, with gene set sizes set to a minimum of 20 and a maximum of 100.

#### ChIP-seq and ATAC-seq data visualisation

The published ChIP-seq dataset (GSE221335) was visualised using the Integrative Genomics Viewer^100^.

### Use of large language models (LLMs)

During the preparation of this work, the authors used ChatGPT Classic (v1.2026.184) to assist with language refinement. After using this tool, the authors reviewed and edited the content as needed and take full responsibility for the content of the publication.

## Data availability

The mass spectrometry proteomics data have been deposited to the ProteomeXchange Consortium via the PRIDE^101^ partner repository with the dataset identifier PXD081575.

## Tables

**Supplementary Table 1:** Variable width isolation windows for mass spectrometry method.

| DIA Window | Min | Max | m/z centre | Window width |
| --- | --- | --- | --- | --- |
| 1 | 350 | 394 | 372.0 | 44 |
| 2 | 393 | 424 | 408.5 | 31 |
| 3 | 423 | 452 | 437.5 | 29 |
| 4 | 451 | 478 | 464.5 | 27 |
| 5 | 477 | 504 | 490.5 | 27 |
| 6 | 503 | 529 | 516.0 | 26 |
| 7 | 528 | 555 | 541.5 | 27 |
| 8 | 554 | 581 | 567.5 | 27 |
| 9 | 580 | 608 | 594.0 | 28 |
| 10 | 607 | 635 | 621.0 | 28 |
| 11 | 634 | 663 | 648.5 | 29 |
| 12 | 662 | 693 | 677.5 | 31 |
| 13 | 692 | 725 | 708.5 | 33 |
| 14 | 724 | 759 | 741.5 | 35 |
| 15 | 758 | 798 | 778.0 | 40 |
| 16 | 797 | 841 | 819.0 | 44 |
| 17 | 840 | 892 | 866.0 | 52 |
| 18 | 891 | 959 | 925.0 | 68 |
| 19 | 958 | 1062 | 1010.0 | 104 |
| 20 | 1061 | 1650 | 1355.5 | 589 |

**Supplementary Table 2:** Antibodies used for immunoblotting.

| Antibodies | Source | Identifier |
| --- | --- | --- |
| ETS1 | Abcam | Cat. #ab220361 |
| Histone H3 | Cell Signaling Technology | Cat. #4499 |
| Phospho-TBC1D4 (Thr642) | Cell Signaling Technology | Cat. #8881 |
| TBC1D4 (rabbit; polyclonal) | Generated in-house | N/A |
| Phospho-AKT (Thr308) | Cell Signaling Technology | Cat. #13038 |
| Phospho-AKT (Ser473) | Cell Signaling Technology | Cat. #4051 |
| Pan-AKT | Cell Signaling Technology | Cat. #4685 |
| Phospho-PRAS40 (Thr246) | Cell Signaling Technology | Cat. #2997 |
| PRAS40 | Cell Signaling Technology | Cat. #2691 |
| Pan 14-3-3 | Santa Cruz | Cat. #sc-1657 |
| PRDX3 | Invitrogen | Cat. #LF-PA0030 |
| α-tubulin | Cell Signaling Technology | Cat. #3873 |
| Anti-TfR (CD71; Rat) | Thermo Fisher Scientific | Cat. #14-0711-82 |
| WGA-FITC lectin | Merck | Cat. #L4895 |
| Alexa Fluor™ 555 (Goat; anti-Rat) | Invitrogen | Cat. #A21434 |

## Acknowledgments

We would like to thank the Sydney Mass Spectrometry facility and Large Animal Services in the Charles Perkins Centre at The University of Sydney for mass spectrometry and mouse housing support. This work was supported by an Australian Research Council Laureate Fellowship (to D.E.J.), an Australian Government Research Training Program Scholarship (to H.B.C., J.Sc., A.L.), and a University of Sydney Postgraduate Award (to Y.J.).

## Author information

These authors contributed equally: Yi Lin Jiang, Kristen C. Cooke.

## Contributions

Conceptualization: D.E.J., G.J.C., J.St., K.C.C. Data curation: Y.J., H.B.C., S.M. Formal analysis: Y.J. Funding acquisition: D.E.J. Investigation: Y.J., K.C.C., H.B.C., M.P., A.D.-V., S.W.C.M., M.E.N., J.Sc., A.L., G.J.C., J.St., D.E.J., S.M. Methodology: K.C.C., J.St., Y.J. Project administration: D.E.J., J.St., K.C.C. Resources: G.M., J.G.B. Software: Y.J. Supervision: D.E.J., S.M., J.St. Validation: K.C.C. Visualization: Y.J. Writing – original draft: Y.J., K.C.C., H.B.C., J.St., D.E.J., S.M. Writing – review & editing: all authors.

## Corresponding authors

Correspondence to Jacqueline Stöckli, David E. James, and Søren Madsen

---

**Extended Data Fig. 1.**
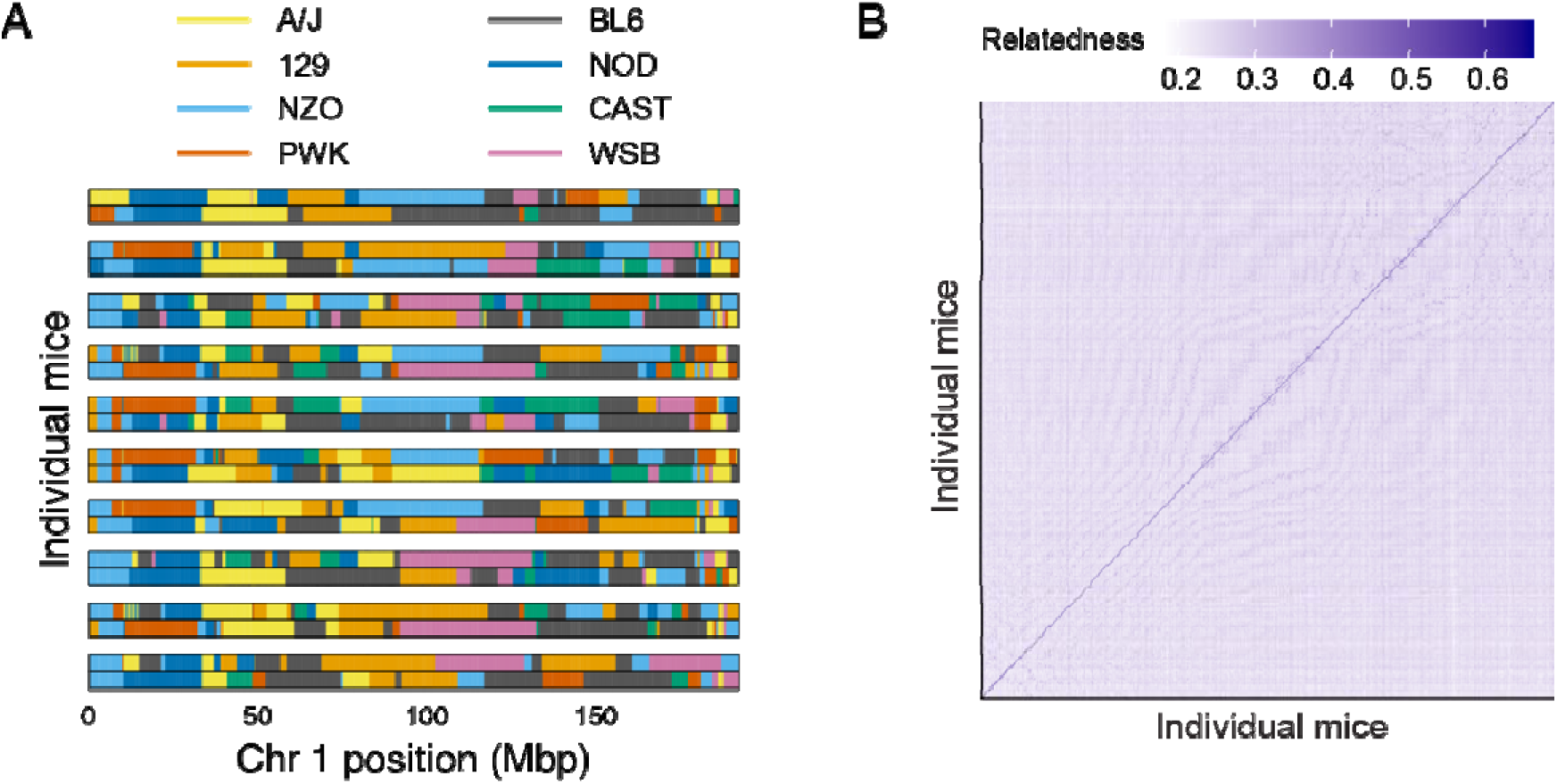
Genetic diversity in the DOz population. **(A)** Founder diplotype composition across chromosome 1 in 10 randomly selected DOz mice. **(B)** Kinship analysis of the DOz mice showing low genetic relatedness.

**Extended Data Fig. 2.**
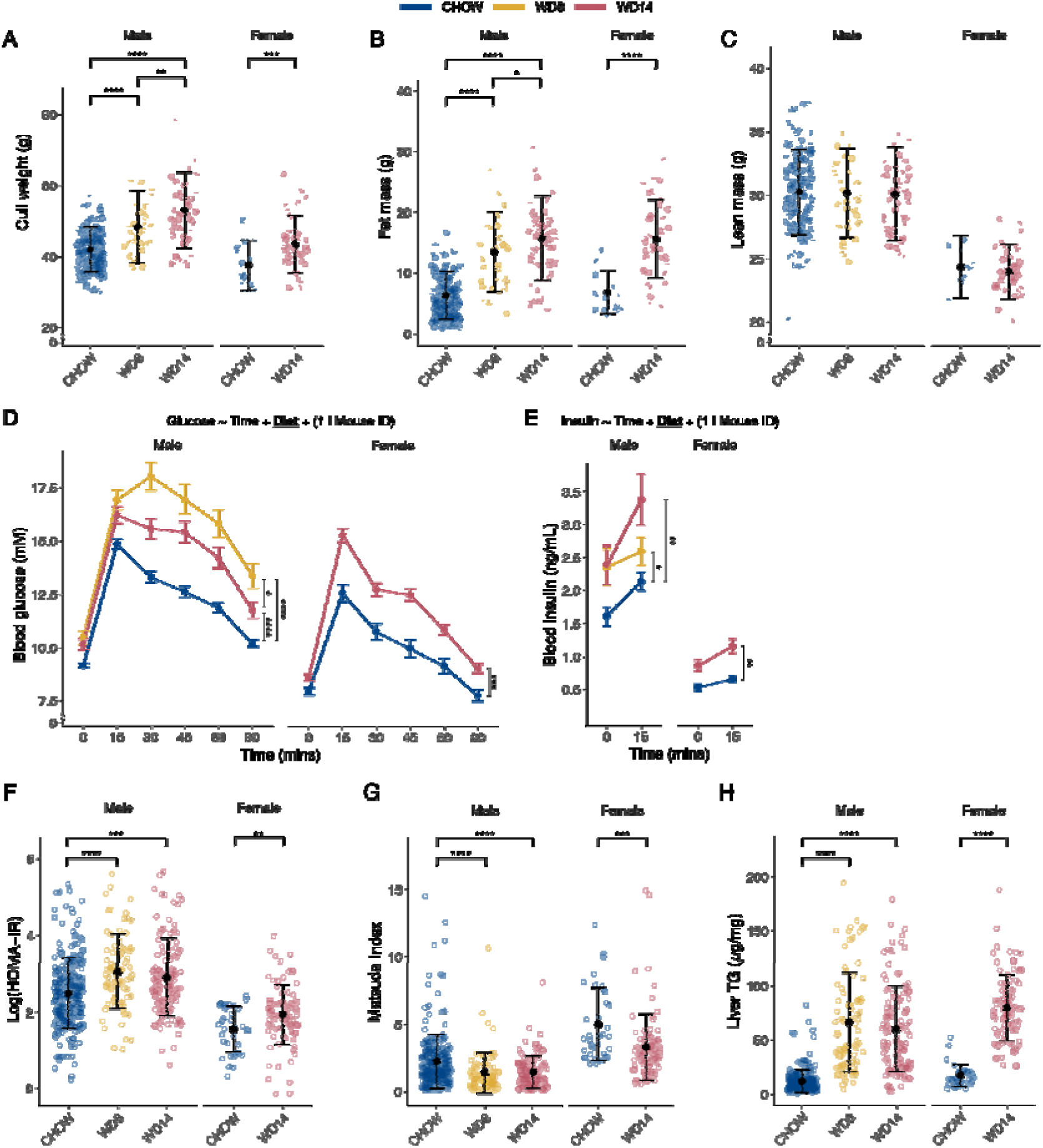
Systemic metabolic measures of DOz mice under different dietary interventions. **(A)** Cull weight. **(B)** Fat mass. **(C)** Lean mass. **(D)** Blood glucose and **(E)** insulin concentrations during an oral GTT. **(F-G)** Systemic insulin resistance/sensitivity indices calculated based on glucose tolerance test (GTT) results: Log (HOMA-IR) (F) and Matsuda Index (G). **(H)** Liver triglyceride (TG) level. For (A-C, F-H), data are presented as mean ± SD. For (D-E), data are presented as mean ± SE. For (A-C, F), t-tests were performed between groups; Wilcoxon tests were performed for (G-H). For each sex in (D, E), glucose and insulin time courses were analysed using linear mixed-effects models with diet and time as fixed effects and mouse ID as a random effect. Pairwise diet comparisons were performed using the same model, with Benjamini–Hochberg correction for multiple testing. Asterisks denote the statistical significance of the diet effect

**Extended Data Fig. 3.**
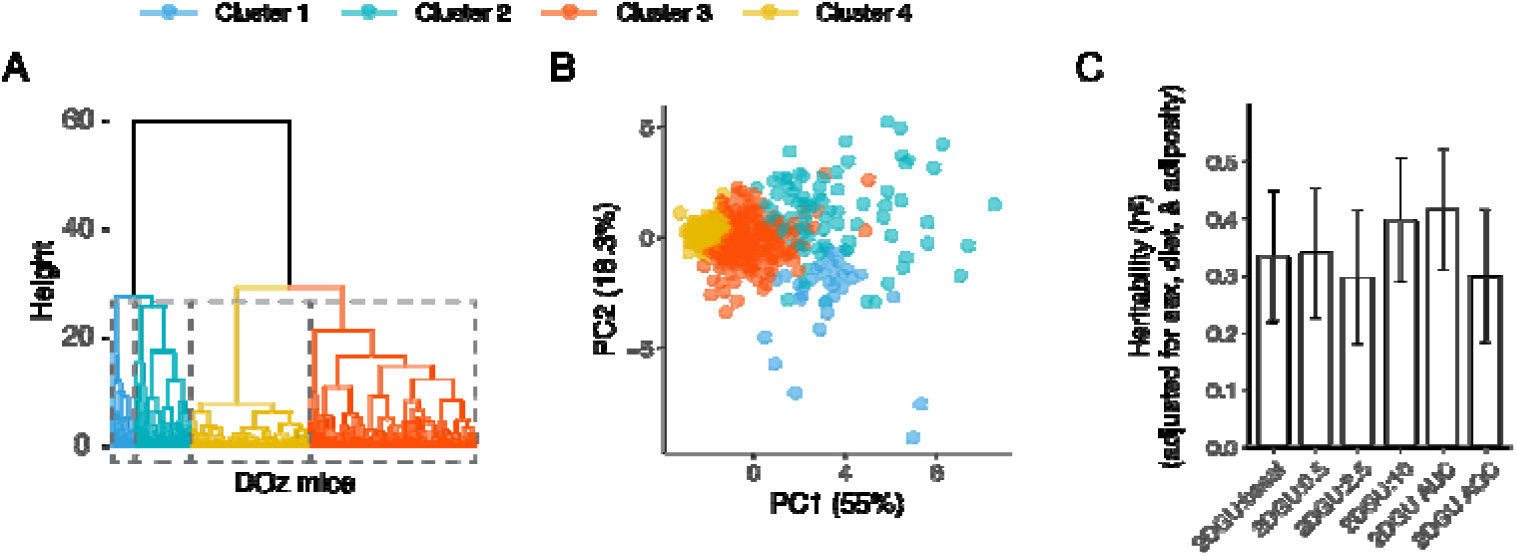
Clustering and heritability of the 2DGU traits. **(A)** Dendrogram showing the hierarchical clustering, using both absolute 2DGU values across insulin doses and their derived pairwise differences, along with the **(B)** associated PCA, coloured by clusters. Percentages of variance explained by PC1 and PC2 are indicated on the axes. **(C)** Heritability of 2DGU measures in the DOz population, adjusting for sex, diet, and adiposity. Error bars are heritability ± SE.

**Extended Data Fig. 4.**
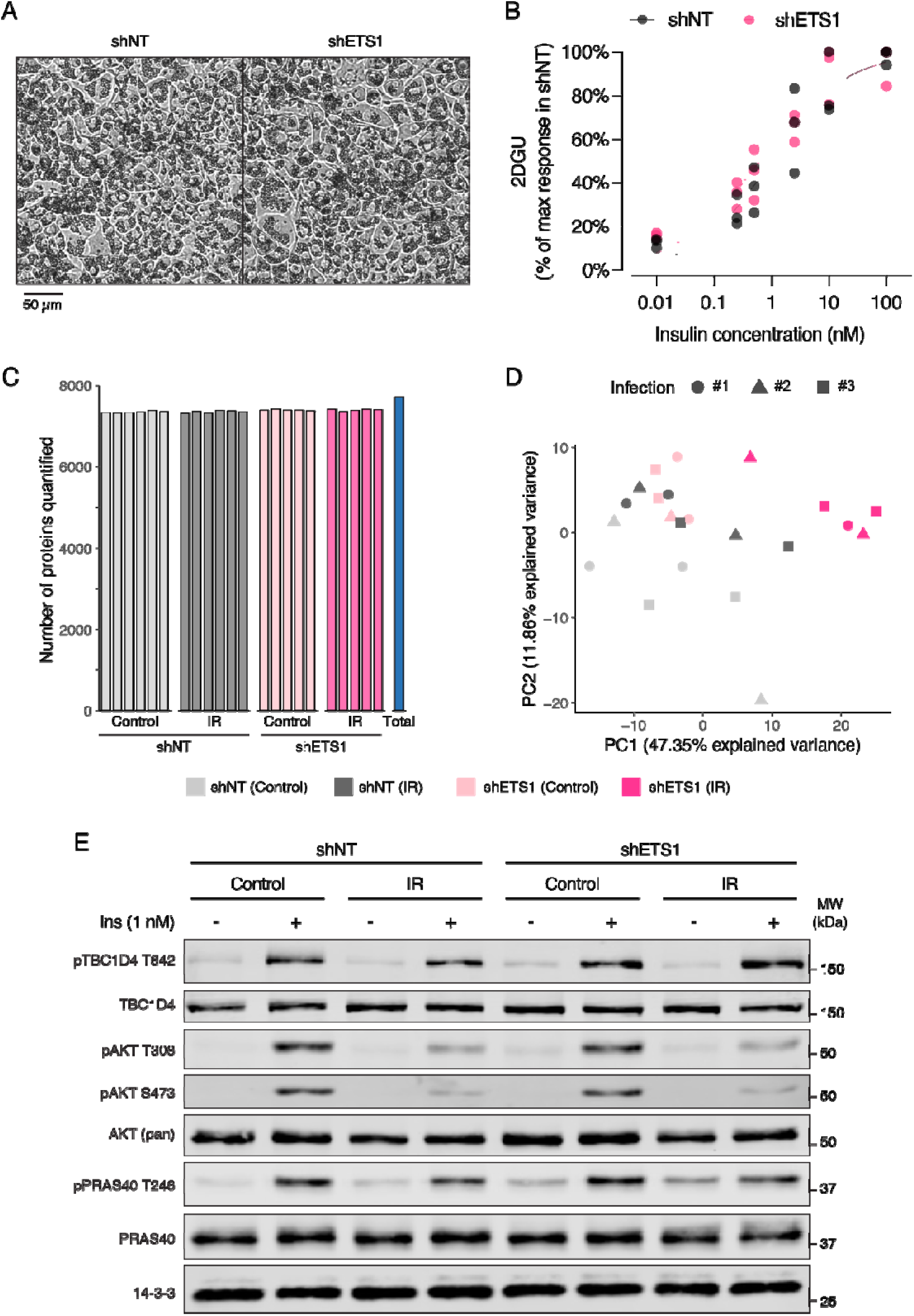
Additional cellular characterisations and proteomic analyses of ETS1 KD in 3T3-L1 adipocytes under control or insulin-resistant (IR) states. **(A)** Representative brightfield images of differentiated shNT or shETS1 adipocytes at the control state. Scale bar is 50 μm. **(B)** 2DGU was measured in shNT or shETS1 adipocytes under control conditions at different insulin doses. Individual data points represent biological replicates. **(C)** Number of proteins quantified for each biological replicate within each condition group. **(D)** PCA of shNT and shETS1 adipocytes under control and IR states across three independent retroviral infections. Data points represent biological replicates, coloured by experimental group and shaped by infection batch. Percentages of variance explained by PC1 and PC2 are indicated on the axes. **(E)** shNT and shETS1 adipocytes were maintained under control or IR conditions and stimulated with or without 1 nM insulin for 10 mins. Cell lysates were immunoblotted, and expression and phosphorylation of AS160, AKT, and PRAS40 were assessed as readouts of canonical insulin signalling. 14-3-3 was used as a loading control. Representative blots are shown.

**Extended Data Fig. 5.**
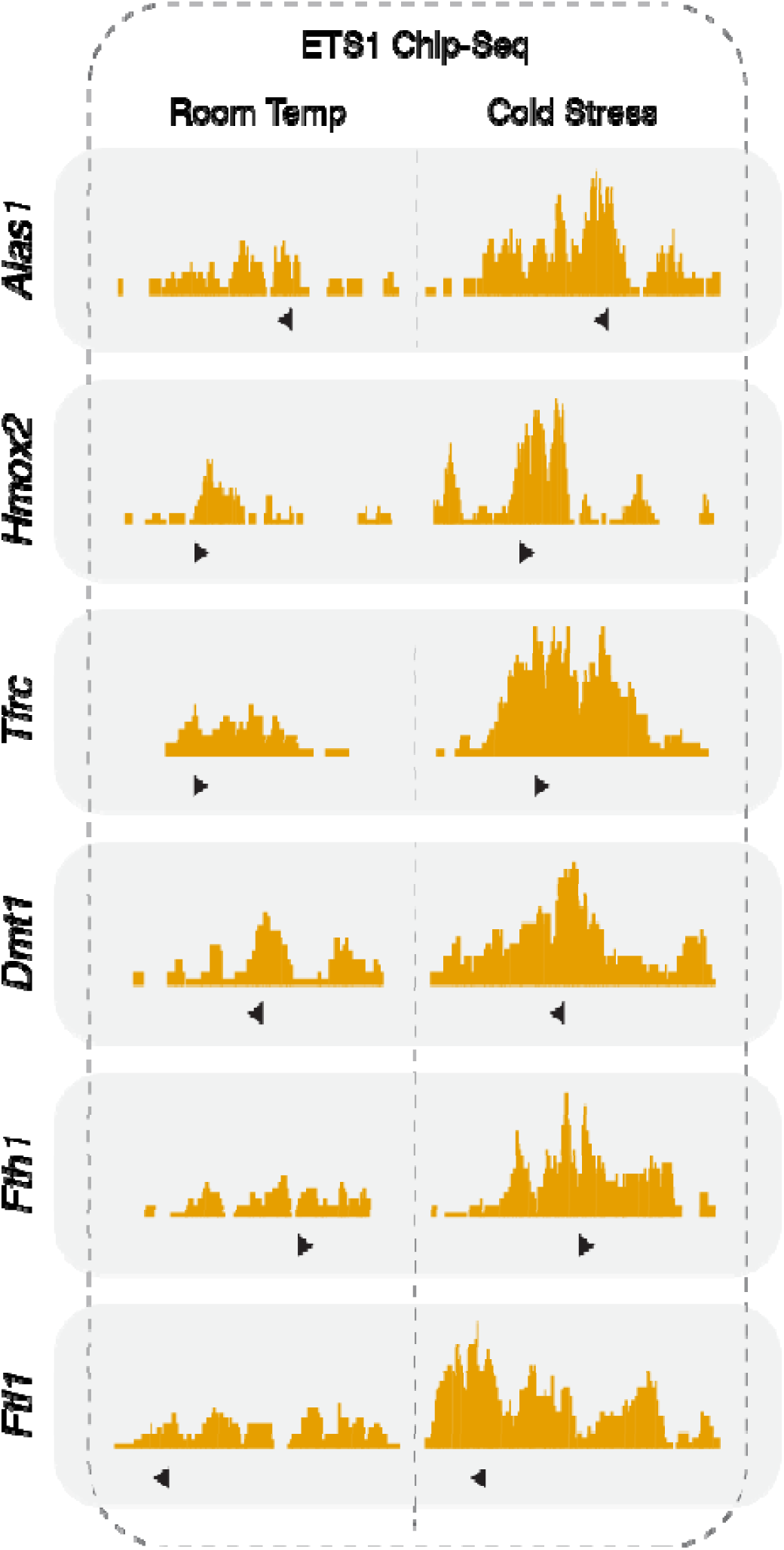
ETS1 binding at the promoter regions of heme- and iron-related genes. The Integrative Genomics Viewer was used to visualise ETS1 binding at the promoter regions of *Alas1*, *Hmox2*, *Tfrc*, *Dmt1* (also known as *Slc11a2*), *Fth1*, and *Ftl1*, using published ChIP-seq datasets (GSE221335). ETS1 ChIP-seq tracks from inguinal WAT under room temperature and cold stress conditions were included to illustrate the stress-responsive binding of ETS1 to these target genes. Triangles indicate the direction of the transcription start site.

**Supplementary Fig. 1.**
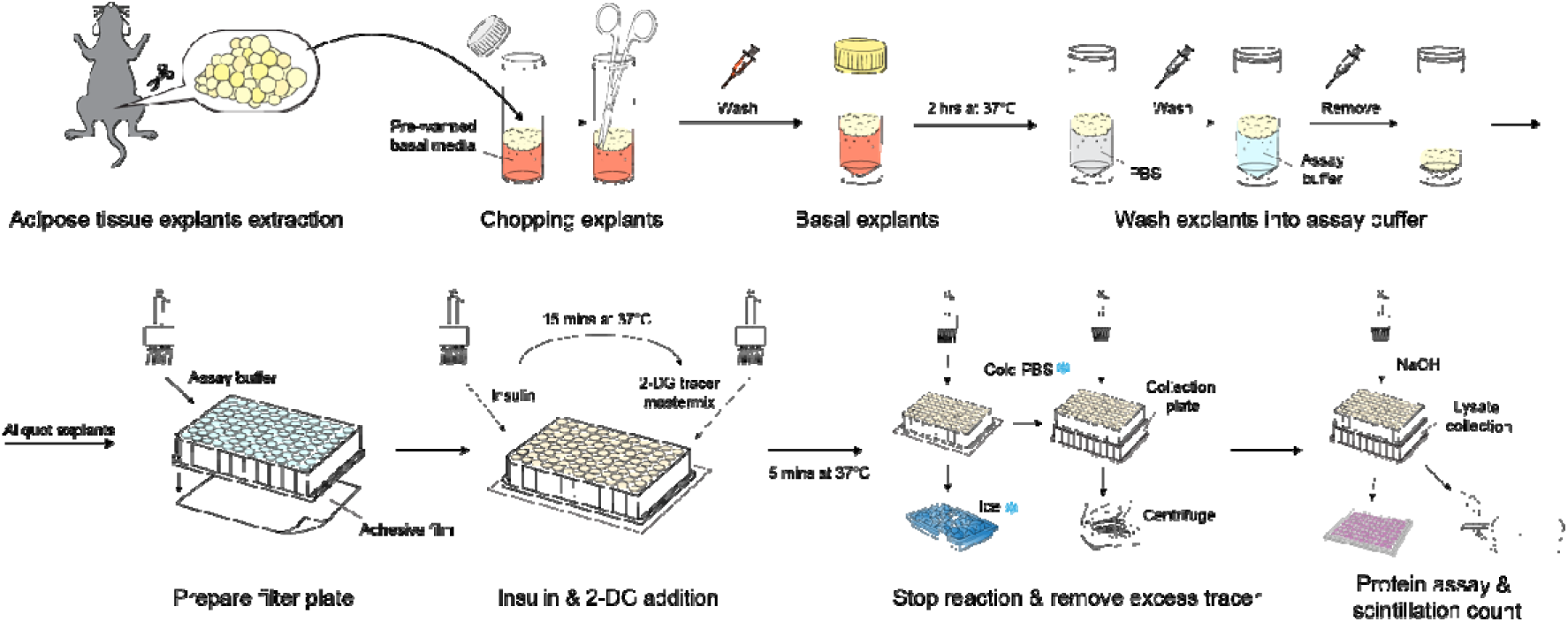
Procedure of the high-throughput *ex vivo* 2DGU assay.

## References

1 Harvey, I., Boudreau, A. & Stephens, J. M. Adipose tissue in health and disease. Open Biology 10, 200291 (2020). 10.1098/rsob.200291

2 James, D. E., Stöckli, J. & Birnbaum, M. J. The aetiology and molecular landscape of insulin resistance. Nature Reviews Molecular Cell Biology 22, 751–771 (2021). 10.1038/s41580-021-00390-6

3 Scheja, L. & Heeren, J. The endocrine function of adipose tissues in health and cardiometabolic disease. Nature Reviews Endocrinology 15, 507–524 (2019). 10.1038/s41574-019-0230-6

4 Kershaw, E. E. & Flier, J. S. Adipose tissue as an endocrine organ. J Clin Endocrinol Metab 89, 2548–2556 (2004). 10.1210/jc.2004-0395

5 Santoro, A., McGraw, T. E. & Kahn, B. B. Insulin action in adipocytes, adipose remodeling, and systemic effects. Cell Metabolism 33, 748–757 (2021). 10.1016/j.cmet.2021.03.019

6 James, D. E., Brown, R., Navarro, J. & Pilch, P. F. Insulin-regulatable tissues express a unique insulin-sensitive glucose transport protein. Nature 333, 183–185 (1988). 10.1038/333183a0

7 Rosen, E. D. & Spiegelman, B. M. Adipocytes as regulators of energy balance and glucose homeostasis. Nature 444, 847–853 (2006). 10.1038/nature05483

8 Krycer, J. R. et al. Lactate production is a prioritized feature of adipocyte metabolism. J Biol Chem 295, 83–98 (2020). 10.1074/jbc.RA119.011178

9 Abel, E. D. et al. Adipose-selective targeting of the GLUT4 gene impairs insulin action in muscle and liver. Nature 409, 729–733 (2001). 10.1038/35055575

10 Chen, J. et al. The trans-ancestral genomic architecture of glycemic traits. Nature Genetics 53, 840–860 (2021). 10.1038/s41588-021-00852-9

11 Manning, A. K. et al. A genome-wide approach accounting for body mass index identifies genetic variants influencing fasting glycemic traits and insulin resistance. Nat Genet 44, 659–669 (2012). 10.1038/ng.2274

12 Williamson, A. et al. Genome-wide association study and functional characterization identifies candidate genes for insulin-stimulated glucose uptake. Nature Genetics 55, 973–983 (2023). 10.1038/s41588-023-01408-9

13 Blüher, M. Metabolically Healthy Obesity. Endocrine Reviews 41, bnaa004 (2020). 10.1210/endrev/bnaa004

14 Schulze, M. B. & Stefan, N. Metabolically healthy obesity: from epidemiology and mechanisms to clinical implications. Nature Reviews Endocrinology 20, 633–646 (2024). 10.1038/s41574-024-01008-5

15 Ghaben, A. L. & Scherer, P. E. Adipogenesis and metabolic health. Nature Reviews Molecular Cell Biology 20, 242–258 (2019). 10.1038/s41580-018-0093-z

16 Burhans, M. S., Hagman, D. K., Kuzma, J. N., Schmidt, K. A. & Kratz, M. Contribution of Adipose Tissue Inflammation to the Development of Type 2 Diabetes Mellitus. Compr Physiol 9, 1–58 (2018). 10.1002/cphy.c170040

17 Maniyadath, B., Zhang, Q., Gupta, R. K. & Mandrup, S. Adipose tissue at single-cell resolution. Cell Metabolism 35, 386–413 (2023). 10.1016/j.cmet.2023.02.002

18 Smith, G. I., Mittendorfer, B. & Klein, S. Metabolically healthy obesity: facts and fantasies. J Clin Invest 129, 3978–3989 (2019). 10.1172/jci129186

19 Nelson, M. E. et al. Systems-level analysis of insulin action in mouse strains provides insight into tissue- and pathway-specific interactions that drive insulin resistance. Cell Metabolism 34, 227–239.e226 (2022). 10.1016/j.cmet.2021.12.013

20 Masson, S. W. C. et al. Leveraging genetic diversity to identify small molecules that reverse mouse skeletal muscle insulin resistance. eLife 12, RP86961 (2023). 10.7554/eLife.86961

21 Ferrannini, E. et al. Adipose tissue and skeletal muscle insulin-mediated glucose uptake in insulin resistance: role of blood flow and diabetes. Am J Clin Nutr 108, 749–758 (2018). 10.1093/ajcn/nqy162

22 Minard, Annabel Y. et al. mTORC1 Is a Major Regulatory Node in the FGF21 Signaling Network in Adipocytes. Cell Reports 17, 29–36 (2016). 10.1016/j.celrep.2016.08.086

23 Muise, E. S. et al. Downstream Signaling Pathways in Mouse Adipose Tissues Following Acute In Vivo Administration of Fibroblast Growth Factor 21. PLOS ONE 8, e73011 (2013). 10.1371/journal.pone.0073011

24 Mullins, G. R. et al. Catecholamine-induced lipolysis causes mTOR complex dissociation and inhibits glucose uptake in adipocytes. Proc Natl Acad Sci U S A 111, 17450–17455 (2014). 10.1073/pnas.1410530111

25 Hue, L. & Taegtmeyer, H. The Randle cycle revisited: a new head for an old hat. Am J Physiol Endocrinol Metab 297, E578–591 (2009). 10.1152/ajpendo.00093.2009

26 Randle, P. J., Garland, P. B., Hales, C. N. & Newsholme, E. A. The glucose fatty-acid cycle. Its role in insulin sensitivity and the metabolic disturbances of diabetes mellitus. Lancet 1, 785–789 (1963). 10.1016/s0140-6736(63)91500-9

27 Wang, Q. A., Tao, C., Gupta, R. K. & Scherer, P. E. Tracking adipogenesis during white adipose tissue development, expansion and regeneration. Nature Medicine 19, 1338–1344 (2013). 10.1038/nm.3324

28 Jeffery, E., Church, C. D., Holtrup, B., Colman, L. & Rodeheffer, M. S. Rapid depot-specific activation of adipocyte precursor cells at the onset of obesity. Nat Cell Biol 17, 376–385 (2015). 10.1038/ncb3122

29 Loos, R. J. F. & Yeo, G. S. H. The genetics of obesity: from discovery to biology. Nature Reviews Genetics 23, 120–133 (2022). 10.1038/s41576-021-00414-z

30 Elks, C. E. et al. Variability in the Heritability of Body Mass Index: A Systematic Review and Meta-Regression. Frontiers in Endocrinology Volume 3 - 2012 (2012). 10.3389/fendo.2012.00029

31 Broman, K. W. et al. R/qtl2: Software for Mapping Quantitative Trait Loci with High-Dimensional Data and Multiparent Populations. Genetics 211, 495–502 (2019). 10.1534/genetics.118.301595

32 Ruby, M. A., Riedl, I., Massart, J., Åhlin, M. & Zierath, J. R. Protein kinase N2 regulates AMP kinase signaling and insulin responsiveness of glucose metabolism in skeletal muscle. American Journal of Physiology-Endocrinology and Metabolism 313, E483–E491 (2017). 10.1152/ajpendo.00147.2017

33 Sun, Y. et al. Lmo4-resistin signaling contributes to adipose tissue-liver crosstalk upon weight cycling. The FASEB Journal 34, 4732–4748 (2020). 10.1096/fj.201902708R

34 Rogers, S. et al. Identification of a novel glucose transporter-like protein—GLUT-12. American Journal of Physiology-Endocrinology and Metabolism 282, E733–E738 (2002). 10.1152/ajpendo.2002.282.3.E733

35 Stuart, C. A., Howell, M. E. A., Zhang, Y. & Yin, D. Insulin-Stimulated Translocation of Glucose Transporter (GLUT) 12 Parallels That of GLUT4 in Normal Muscle. The Journal of Clinical Endocrinology & Metabolism 94, 3535–3542 (2009). 10.1210/jc.2009-0162

36 Sancar, G. & Birkenfeld, A. L. The role of adipose tissue dysfunction in hepatic insulin resistance and T2D. Journal of Endocrinology 262, e240115 (2024). 10.1530/JOE-24-0115

37 Cutler, H. B. et al. Synteny – a high throughput web tool to streamline causal gene prioritisation and provide insight into protein function. Scientific Reports 15, 44761 (2025). 10.1038/s41598-025-28473-w

38 Diaz-Vegas, A. et al. A high-content endogenous GLUT4 trafficking assay reveals new aspects of adipocyte biology. Life Science Alliance 6, e202201585 (2023). 10.26508/lsa.202201585

39 Fazakerley, D. J. et al. Phosphoproteomics reveals rewiring of the insulin signaling network and multi-nodal defects in insulin resistance. Nature Communications 14, 923 (2023). 10.1038/s41467-023-36549-2

40 Wang, H. et al. Sex-specific phosphoproteome responses to calorie restriction and insulin in skeletal muscle from older rats. The Journals of Gerontology: Series A 80, glaf231 (2025). 10.1093/gerona/glaf231

41 Dutt, S., Hamza, I. & Bartnikas, T. B. Molecular Mechanisms of Iron and Heme Metabolism. Annual Review of Nutrition 42, 311–335 (2022). 10.1146/annurev-nutr-062320-112625

42 Kakhlon, O. & Cabantchik, Z. I. The labile iron pool: characterization, measurement, and participation in cellular processes. Free Radical Biology and Medicine 33, 1037–1046 (2002). 10.1016/S0891-5849(02)01006-7

43 Tanner, L. I. & Lienhard, G. E. Insulin elicits a redistribution of transferrin receptors in 3T3-L1 adipocytes through an increase in the rate constant for receptor externalization. Journal of Biological Chemistry 262, 8975–8980 (1987). 10.1016/S0021-9258(18)48032-5

44 Davis, R. J., Corvera, S. & Czech, M. P. Insulin stimulates cellular iron uptake and causes the redistribution of intracellular transferrin receptors to the plasma membrane. Journal of Biological Chemistry 261, 8708–8711 (1986). 10.1016/S0021-9258(19)84438-1

45 Orino, K. et al. Ferritin and the response to oxidative stress. Biochemical Journal 357, 241–247 (2001). 10.1042/bj3570241

46 Hoehn, K. L. et al. Insulin resistance is a cellular antioxidant defense mechanism. Proceedings of the National Academy of Sciences 106, 17787–17792 (2009). 10.1073/pnas.0902380106

47 Fazakerley, D. J. et al. Mitochondrial CoQ deficiency is a common driver of mitochondrial oxidants and insulin resistance. eLife 7, e32111 (2018). 10.7554/eLife.32111

48 Fazakerley, D. J. et al. Mitochondrial oxidative stress causes insulin resistance without disrupting oxidative phosphorylation. J Biol Chem 293, 7315–7328 (2018). 10.1074/jbc.RA117.001254

49 Wu, S. et al. M2 macrophages independently promote beige adipogenesis via blocking adipocyte Ets1. Nature Communications 15, 1646 (2024). 10.1038/s41467-024-45899-4

50 Dittmer, J. The biology of the Ets1 proto-oncogene. Mol Cancer 2, 29 (2003). 10.1186/1476-4598-2-29

51 Guilherme, A., Virbasius, J. V., Puri, V. & Czech, M. P. Adipocyte dysfunctions linking obesity to insulin resistance and type 2 diabetes. Nat Rev Mol Cell Biol 9, 367–377 (2008). 10.1038/nrm2391

52 Agrawal, S. et al. BMI-adjusted adipose tissue volumes exhibit depot-specific and divergent associations with cardiometabolic diseases. Nature Communications 14, 266 (2023). 10.1038/s41467-022-35704-5

53 Shungin, D. et al. New genetic loci link adipose and insulin biology to body fat distribution. Nature 518, 187–196 (2015). 10.1038/nature14132

54 Locke, A. E. et al. Genetic studies of body mass index yield new insights for obesity biology. Nature 518, 197–206 (2015). 10.1038/nature14177

55 Morgan, D. et al. The transcription complex p52–ETS1 is essential for germinal center formation. Nature Immunology 26, 1553–1566 (2025). 10.1038/s41590-025-02236-1

56 Yan, M. et al. ETS1 governs pathological tissue-remodeling programs in disease-associated fibroblasts. Nature Immunology 23, 1330–1341 (2022). 10.1038/s41590-022-01285-0

57 Chandra, A. et al. Quantitative control of *Ets1* dosage by a multi-enhancer hub promotes Th1 cell differentiation and protects from allergic inflammation. Immunity 56, 1451–1467.e1412 (2023). 10.1016/j.immuni.2023.05.004

58 Yang, M. et al. Role of the E26 transformation specific transcription factor family in metabolic disorders. Journal of Endocrinological Investigation 48, 2547–2561 (2025). 10.1007/s40618-025-02634-0

59 Chen, F. et al. Transcription factor Ets-1 links glucotoxicity to pancreatic beta cell dysfunction through inhibiting PDX-1 expression in rodent models. Diabetologia 59, 316–324 (2016). 10.1007/s00125-015-3805-3

60 Luo, Y. et al. Transcription Factor Ets1 Regulates Expression of Thioredoxin-Interacting Protein and Inhibits Insulin Secretion in Pancreatic β-Cells. PLOS ONE 9, e99049 (2014). 10.1371/journal.pone.0099049

61 Zhang, X.-F., Zhu, Y., Liang, W.-B. & Zhang, J.-J. Transcription factor Ets-1 inhibits glucose-stimulated insulin secretion of pancreatic β-cells partly through up-regulation of COX-2 gene expression. Endocrine 46, 470–476 (2014). 10.1007/s12020-013-0114-9

62 Li, K. et al. Ets1-Mediated Acetylation of FoxO1 Is Critical for Gluconeogenesis Regulation during Feed-Fast Cycles. Cell Reports 26, 2998–3010.e2995 (2019). 10.1016/j.celrep.2019.02.035

63 Song, J., et al. Ets1-dependent Mek-Erk signaling drives adipose tissue macrophage anti-inflammatory polarization to ameliorates insulin resistance. Cell Reports 45 (2026). 10.1016/j.celrep.2026.117244

64 Zhang, Z. et al. Adipocyte iron levels impinge on a fat-gut crosstalk to regulate intestinal lipid absorption and mediate protection from obesity. Cell Metabolism 33, 1624–1639.e1629 (2021). 10.1016/j.cmet.2021.06.001

65 Hilton, C., Sabaratnam, R., Drakesmith, H. & Karpe, F. Iron, glucose and fat metabolism and obesity: an intertwined relationship. International Journal of Obesity 47, 554–563 (2023). 10.1038/s41366-023-01299-0

66 Cooksey, R. C. et al. Dietary iron restriction or iron chelation protects from diabetes and loss of beta-cell function in the obese (ob/ob lep-/-) mouse. Am J Physiol Endocrinol Metab 298, E1236–1243 (2010). 10.1152/ajpendo.00022.2010

67 Houschyar, K. S. et al. Effects of phlebotomy-induced reduction of body iron stores on metabolic syndrome: results from a randomized clinical trial. BMC Medicine 10, 54 (2012). 10.1186/1741-7015-10-54

68 Yan, H.-F., Liu, Z.-Y., Guan, Z.-A. & Guo, C. Deferoxamine ameliorates adipocyte dysfunction by modulating iron metabolism in ob/ob mice. Endocrine Connections 7, 604–616 (2018). 10.1530/EC-18-0054

69 Barton, J. C. & Acton, R. T. Diabetes in HFE Hemochromatosis. J Diabetes Res 2017, 9826930 (2017). 10.1155/2017/9826930

70 Fernández-Real, J. M. et al. Blood letting in high-ferritin type 2 diabetes: effects on insulin sensitivity and beta-cell function. Diabetes 51, 1000–1004 (2002). 10.2337/diabetes.51.4.1000

71 Valenti, L. et al. Iron depletion by phlebotomy improves insulin resistance in patients with nonalcoholic fatty liver disease and hyperferritinemia: evidence from a case-control study. Am J Gastroenterol 102, 1251–1258 (2007). 10.1111/j.1572-0241.2007.01192.x

72 Gabrielsen, J. S. et al. Adipocyte iron regulates adiponectin and insulin sensitivity. J Clin Invest 122, 3529–3540 (2012). 10.1172/jci44421

73 Wlazlo, N. et al. Iron Metabolism Is Associated With Adipocyte Insulin Resistance and Plasma Adiponectin: The Cohort on Diabetes and Atherosclerosis Maastricht (CODAM) study. Diabetes Care 36, 309–315 (2013). 10.2337/dc12-0505

74 Plotnik, J. P., Budka, J. A., Ferris, M. W. & Hollenhorst, P. C. ETS1 is a genome-wide effector of RAS/ERK signaling in epithelial cells. Nucleic Acids Res 42, 11928–11940 (2014). 10.1093/nar/gku929

75 Paumelle, R. et al. Hepatocyte growth factor/scatter factor activates the ETS1 transcription factor by a RAS-RAF-MEK-ERK signaling pathway. Oncogene 21, 2309–2319 (2002). 10.1038/sj.onc.1205297

76 Yang, B. S. et al. Ras-mediated phosphorylation of a conserved threonine residue enhances the transactivation activities of c-Ets1 and c-Ets2. Mol Cell Biol 16, 538–547 (1996). 10.1128/mcb.16.2.538

77 Moreno-Navarrete, J. M. et al. HMOX1 as a marker of iron excess-induced adipose tissue dysfunction, affecting glucose uptake and respiratory capacity in human adipocytes. Diabetologia 60, 915–926 (2017). 10.1007/s00125-017-4228-0

78 Thillainadesan, S. et al. The metabolic consequences of ‘yo-yo’ dieting are markedly influenced by genetic diversity. International Journal of Obesity 48, 1170–1179 (2024). 10.1038/s41366-024-01542-2

79 Masson, S. W. C. et al. Genetic variance in the murine defensin locus modulates glucose homeostasis. The EMBO Journal 44, 5694–5711 (2025). 10.1038/s44318-025-00555-5

80 Broman, K. W., Gatti, D. M., Svenson, K. L., Sen, Ś. & Churchill, G. A. Cleaning Genotype Data from Diversity Outbred Mice. G3 Genes|Genomes|Genetics 9, 1571–1579 (2019). 10.1534/g3.119.400165

81 Stöckli, J. et al. ABHD15 regulates adipose tissue lipolysis and hepatic lipid accumulation. Molecular Metabolism 25, 83–94 (2019). 10.1016/j.molmet.2019.05.002

82 Todaro, G. J. & Green, H. Quantitative studies of the growth of mouse embryo cells in culture and their development into established lines. J Cell Biol 17, 299–313 (1963). 10.1083/jcb.17.2.299

83 Zhao, J.-y., Osipovich, O., Koues, O. I., Majumder, K. & Oltz, E. M. Activation of Mouse Tcrb: Uncoupling RUNX1 Function from Its Cooperative Binding with ETS1. The Journal of Immunology 199, 1131–1141 (2017). 10.4049/jimmunol.1700146

84 Zhang, Y. et al. Integrated analysis of the adipocyte plasma membrane proteome reveals KCC1 and PIT2 as novel insulin-responsive transporters. Journal of Biological Chemistry 302 (2026). 10.1016/j.jbc.2026.111282

85 Schreiber, E., Matthias, P., Müller, M. M. & Schaffner, W. Rapid detection of octamer binding proteins with ’mini-extracts’, prepared from a small number of cells. Nucleic Acids Res 17, 6419 (1989). 10.1093/nar/17.15.6419

86 Rappsilber, J., Mann, M. & Ishihama, Y. Protocol for micro-purification, enrichment, pre-fractionation and storage of peptides for proteomics using StageTips. Nat Protoc 2, 1896–1906 (2007). 10.1038/nprot.2007.261

87 Hanna, D. A. et al. Heme oxygenase-2 (HO-2) binds and buffers labile ferric heme in human embryonic kidney cells. Journal of Biological Chemistry 298, 101549 (2022). 10.1016/j.jbc.2021.101549

88 Morrison, G. R. Fluorometric Microdetermination of Heme Protein. Analytical Chemistry 37, 1124–1126 (1965). 10.1021/ac60228a014

89 Sinclair, P. R., Gorman, N. & Jacobs, J. M. Measurement of Heme Concentration. Current Protocols in Toxicology 00, 8.3.1–8.3.7 (1999). 10.1002/0471140856.tx0803s00

90 Abbasi, U., Abbina, S., Gill, A., Bhagat, V. & Kizhakkedathu, J. N. A facile colorimetric method for the quantification of labile iron pool and total iron in cells and tissue specimens. Scientific Reports 11, 6008 (2021). 10.1038/s41598-021-85387-z

91 Duarte, T. L. & Neves, J. V. Measurement of Tissue Non-Heme Iron Content using a Bathophenanthroline-Based Colorimetric Assay. J Vis Exp (2022). 10.3791/63469

92 Benjamini, Y. & Hochberg, Y. Controlling the False Discovery Rate: A Practical and Powerful Approach to Multiple Testing. Journal of the Royal Statistical Society: Series B (Methodological) 57, 289–300 (1995). 10.1111/j.2517-6161.1995.tb02031.x

93 Yang, R., Yi, N. & Xu, S. Box–Cox transformation for QTL mapping. Genetica 128, 133–143 (2006). 10.1007/s10709-005-5577-z

94 Matthews, D. R. et al. Homeostasis model assessment: insulin resistance and beta-cell function from fasting plasma glucose and insulin concentrations in man. Diabetologia 28, 412–419 (1985). 10.1007/bf00280883

95 Matsuda, M. & DeFronzo, R. A. Insulin sensitivity indices obtained from oral glucose tolerance testing: comparison with the euglycemic insulin clamp. Diabetes Care 22, 1462–1470 (1999). 10.2337/diacare.22.9.1462

96 snpStats: SnpMatrix and XSnpMatrix classes and methods v. 1.62.0 (2026).

97 Clayton, D. & Leung, H. T. An R package for analysis of whole-genome association studies. Hum Hered 64, 45–51 (2007). 10.1159/000101422

98 Demichev, V., Messner, C. B., Vernardis, S. I., Lilley, K. S. & Ralser, M. DIA-NN: neural networks and interference correction enable deep proteome coverage in high throughput. Nat Methods 17, 41–44 (2020). 10.1038/s41592-019-0638-x

99 Yu, G., Wang, L. G., Han, Y. & He, Q. Y. clusterProfiler: an R package for comparing biological themes among gene clusters. Omics 16, 284–287 (2012). 10.1089/omi.2011.0118

100 Robinson, J. T. et al. Integrative genomics viewer. Nature Biotechnology 29, 24–26 (2011). 10.1038/nbt.1754

101 Perez-Riverol, Y. et al. The PRIDE database at 20 years: 2025 update. Nucleic Acids Res 53, D543–d553 (2025). 10.1093/nar/gkae1011

